# Untangling mechanisms of crude oil toxicity: linking gene expression, morphology and PAHs at two developmental stages in a cold-water fish

**DOI:** 10.1101/2020.09.09.288852

**Authors:** Elin Sørhus, Carey E. Donald, Denis da Silva, Anders Thorsen, Ørjan Karlsen, Sonnich Meier

## Abstract

Early life stages of fish are highly sensitive to crude oil exposure and thus, short term exposures during critical developmental periods could have detrimental consequences for juvenile survival. Here we administered crude oil to Atlantic haddock (*Melanogrammus aeglefinus*) in short term (3-day) exposures at two developmental time periods: before first heartbeat, from gastrulation to cardiac cone stage (*early*), and from first heartbeat to one day before hatching (*late*). A frequent sampling regime enabled us to determine immediate PAH uptake, metabolite formation and gene expression changes. In general, the embryotoxic consequences of an oil exposure were more severe in the *early* exposure animals. Oil droplet fouling in the highest doses resulted in severe cardiac and craniofacial abnormalities. Gene expression changes of Cytochrome 1 a,b,c and d (*cyp1a,b,c,d*), Bone morphogenetic protein 10 (*bmp10*), ABC transporter b1 (*abcb1*) and Rh-associated G-protein (*rhag*) were linked to PAH uptake, occurrence of metabolites of phenanthrene and developmental and functional abnormalities. We detected circulation-independent, oil-induced gene expression changes and separated phenotypes linked to proliferation, growth and disruption of formation events at early and late developmental stages. Our study gives an increased knowledge about developmentally dependent effects of crude oil toxicity. Thus, providing more knowledge and detail to new and several existing adverse outcome pathways of crude oil toxicity.

**Graphical abstract:** 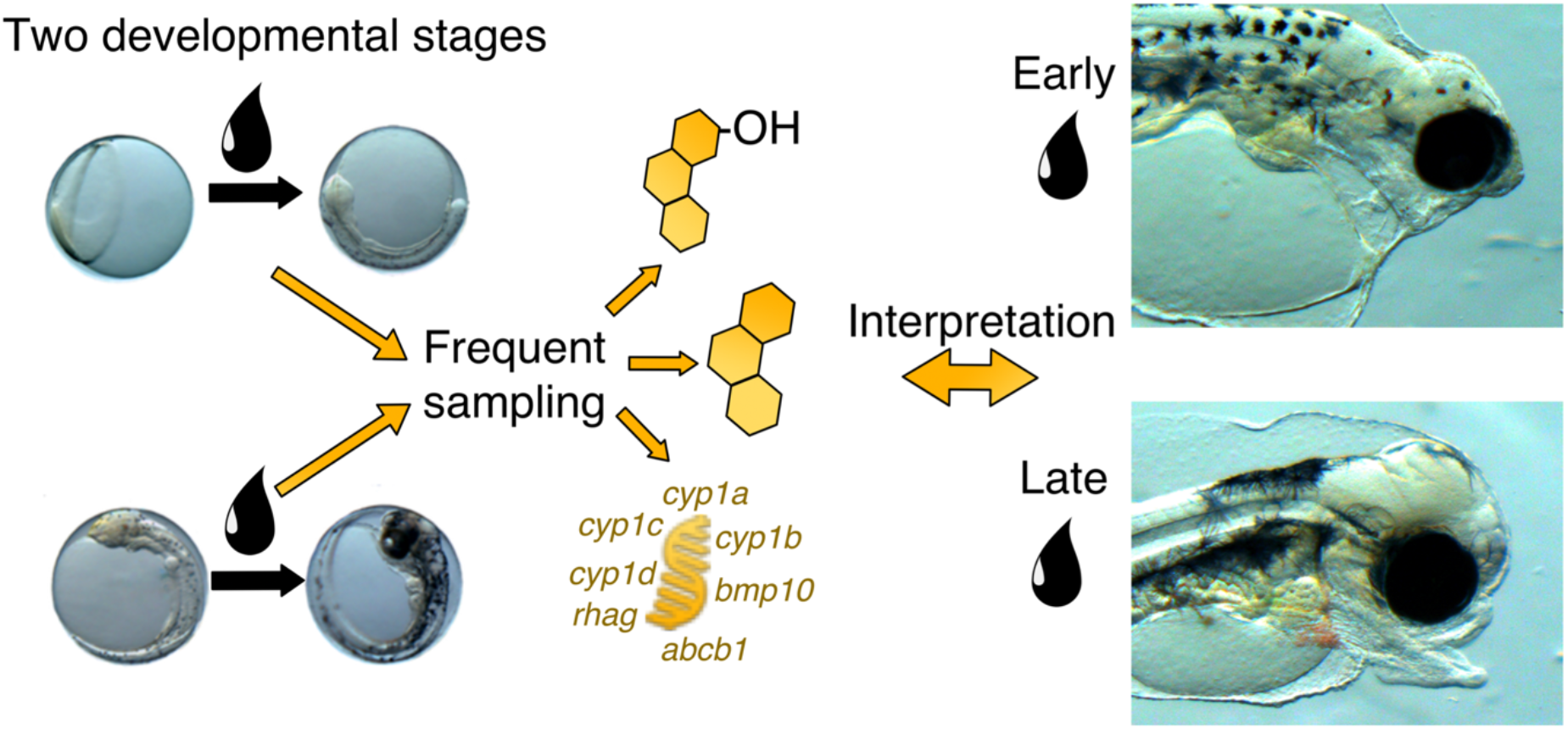

**Highlights:**

- Oil droplet fouling occurred in the whole water column and increased the oil toxicity.
- Early exposure resulted in higher PAH uptake due to lower metabolism resulting in more severe abnormalities.
- A rapid and circulation-indepenent regulation of *bmp10* suggested a direct oil-induced effect on calcium homeostasis.
- Expression of *rhag* indicated a direct oil-induced effect on osmoregulatory cells and osmoregulation.
- Severe eye abnormalities especially in the late exposure was linked to inappropriate overexpression of *cyp1b* in the eyes.

## 1. Introduction

Globally, fish habitats are frequently impacted by accidental oil spills. For fish, consequences of crude oil toxicity are stage dependent, and early life stages (eggs and larvae) are thought to be highly vulnerable for several reasons. Early life stages drift and have no opportunity to avoid contaminated areas compared to juvenile and mature fish (Carroll et al., 2018; Olsen et al., 2010). Relative content of lipids and surface-area-to-volume ratio are higher in early life stages compared to juveniles/adults, making them more susceptible to accumulation of lipophilic environmental toxicants (Petersen and Kristensen, 1998). Finally, critical developmental processes like patterning and organogenesis can be disrupted by crude oil exposure (Incardona, 2017; Pasparakis et al., 2019; Sørhus et al., 2017). However, there are large interspecies sensitivity differences, e.g. the Atlantic haddock (*Melanogrammus aeglefinus*) embryo is approximately 10 times more sensitive to crude oil than its close relative Atlantic cod (Gadus morhua) due to oil droplet fouling on the haddock chorion (Sørensen et al., 2017).

Crude oil is a complex mixture of thousands of different components, however, the content of polycyclic aromatic hydrocarbons (PAHs) appears to correlate to the toxicity in developing fish (Adams et al., 2014; Carls and Meador, 2009). Deactivation and excretion of the larger PAHs (4-6 rings) is initiated by activation of the aryl hydrocarbon receptor (AhR), which in turn induces PAH-degrading enzymes, such as cytochrome P4501A (Cyp1a). Cyp1a induction has for a long time been used as a sensitive biomarker for crude oil toxicity (Goksøyr, 1995). The detoxification process is divided into three phases, with activation of substrate in phase I by *e.g*. cyp1a, followed by conjugation in phase II, and finally transport and excretion in phase III (Wells et al., 2009). Although PAHs are themselves toxic, reactive intermediates (reactive oxygen species, ROS) created during the detoxification process can cause toxicity such as oxidative damage to cellular macromolecules and disruption of signal transduction (Wells et al., 1997).

Cardiac morphology abnormalities can be a result of AhR-dependent mechanisms (Wells et al., 1997), circulation abnormalities (Andres-Delgado and Mercader, 2016) and direct disruption of signaling pathways, i.e calcium (Ebert et al., 2005; Rottbauer et al., 2001; Sørhus et al., 2017). In addition, tricyclic PAHs are shown to cause cardiotoxicity through an AhR-independent manner (Incardona et al., 2004; Marris et al., 2019). Tricyclic PAHs are poor inducers of AhR, and instead act by blocking the repolarizing potassium efflux and a reduction in intracellular calcium cycling (Brette et al., 2014; Brette et al., 2017). The disruption of ionic signaling leads to defects in cardiac rhythm and contractility (Incardona, 2014; Incardona et al., 2009; Sørhus et al., 2016b) as well as calcium regulated cardiomyocyte proliferation (Ebert et al., 2005; Huang et al., 2012; Rottbauer et al., 2001; Sørhus et al., 2017). Beyond these circulatory defects that affect cardiac development (Andres-Delgado and Mercader, 2016), other extra-cardiac abnormalities can be secondary to circulatory defects, including craniofacial and eye development, and osmoregulatory and lipid metabolism deficiencies (Incardona et al., 2004; Incardona and Scholz, 2016; Laurel et al., 2019; Sørhus et al., 2017; Sun et al., 2019)

Numerous other pathways/endpoints have also been reported beyond the effects downstream of circulatory defects described above. PAH-related disruption of retinoid signaling interrupts eye development (Lie et al., 2019). Craniofacial development and growth can be disrupted by inappropriate signaling in formation processes like midline signaling (Abramyan, 2019; Delling et al., 2013; Kimmel et al., 2001) or secondary to impact on craniofacial muscle apparatus (Shwartz et al., 2012). Severe oil-induced aberrations lead to early larval mortality, yet even the smaller developmental impacts can affect survival later in life (Heintz et al., 2000; Hicken et al., 2011). Recently, a study linked oil-induced changes in lipid composition and dynamics to reduced growth and survival in polar cod (*Boreogadus saida*) (Laurel et al., 2019). Consistently, impacts on lipid metabolism pathways have been reported in several RNAseq studies (Sørhus et al., 2017; Xu et al., 2017). Several gene expression studies have started to unravel initiating events and key events of adverse outcome pathways of crude oil (Laurel et al., 2019; Lie et al., 2019; McGruer et al., 2019; Sørhus et al., 2017), but there are still knowledge gaps in understanding the underlying mechanisms of crude oil toxicity.

In this study we aim to improve our understanding of cause and effect relationships by linking immediate transcriptional changes to morphological phenotypes, PAH uptake and metabolism. A frequent and thorough sampling regime during and after exposure enabled us to detect immediate changes. We focused specifically on genes found interesting in a previous study (Sørhus et al., 2017) that are involved in various parts of development and function: cardiac development and calcium homeostasis, osmoregulation and ammonium waste excretion, transport of xenobiotic products, and enzymes possibly involved in phase I xenobiotic metabolism.

This study was designed to understand key events of adverse outcome pathways. First, exposure at the early embryonic stage prior to first heartbeat (2.5-5.5 days post-fertilization (dpf)) allowed us to detect circulation-independent, oil-induced gene expression changes. Second, exposure both during and after completion of organogenesis (7.5-10.5 dpf) enabled us to separate phenotypes linked to proliferation, growth and disruption of formation events at early and late developmental stages. Finally, increased knowledge about uptake, induction and rate of PAH metabolism helps to pinpoint sources of crude oil toxicity. A greater understanding of crude oil toxicity in the framework of adverse outcome pathways is essential for predicting more accurately both acute and delayed consequences of accidental and operational oil spills.

## 2. Materials and methods

### 2.1. Animal collection, maintenance and exposure set up

Fertilized Atlantic haddock (*Melanogrammus aeglefinus*) eggs were collected from tanks with wild-caught broodstock haddock maintained at the Institute of Marine Research, Austevoll Research station. The fertilized eggs were kept in indoor incubators (8±1°C), until transfer of ≈5000 eggs to 50 L green polytethylene plastic experimental tanks. Flow through in the tanks was set to 32 L/hr, the water temperature 8.0°C, and light regime was 12 hours light, 12 hours dark provided by broad spectrum 2×36W Osram Biolux 965 (Osram GmbH, Munich, Germany).

The crude oil used was a weathered blend from the Heidrun oil field of the Norwegian sea. The crude oil is representative of the oil found in Lofoten island areas, an area of both haddock spawning and potential oil industry development. The exposure regime was identical to previous studies (Sørhus et al., 2016b).

Two short (72 hours) exposure experiments were performed, hereafter referred to as *early* and *late* exposure (Fig. 1A). The *early* exposure started at 2.5 dpf and ended at 5.5 dpf (10% epiboly to 10-20 somite/cardiac cone stage), while the *late* exposure started at 7.5 dpf and ended at 10.5 dpf (30-40 somite stage to hatching gland stage). The experimental setup consisted of a control and three exposure doses in a total of 18 tanks (Fig. 1B): control (*C*) 0.0 μg oil/L; low (*L*) 30 μg oil/L; medium (*M*) 100 μg oil/L; and high 300 μg oil/L. In our previous oil exposure experiments with haddock embryos, eggs were allowed to float freely (Sørensen et al., 2017; Sørhus et al., 2015; Sørhus et al., 2016b). Haddock eggs have a natural high buoyancy which allocates them close to the surface, and are potentially exposed to any slick formation. Therefore, eggs were added to plankton mesh chambers (Fig. 1B) in four replicate tanks of each treatment and submerged into the water column (*sub*). To identify an effect of oil droplet binding at the surface, the high dose was administered in three ways: like the other doses, where the eggs were held submerged under the water surface (*H sub*); where the eggs were allowed to float at the surface (*H surf*); and where the eggs were allowed to float and the exposure water was filtered to remove oil microdroplets (water soluble fraction; *H WSF*).

**Figure 1:**
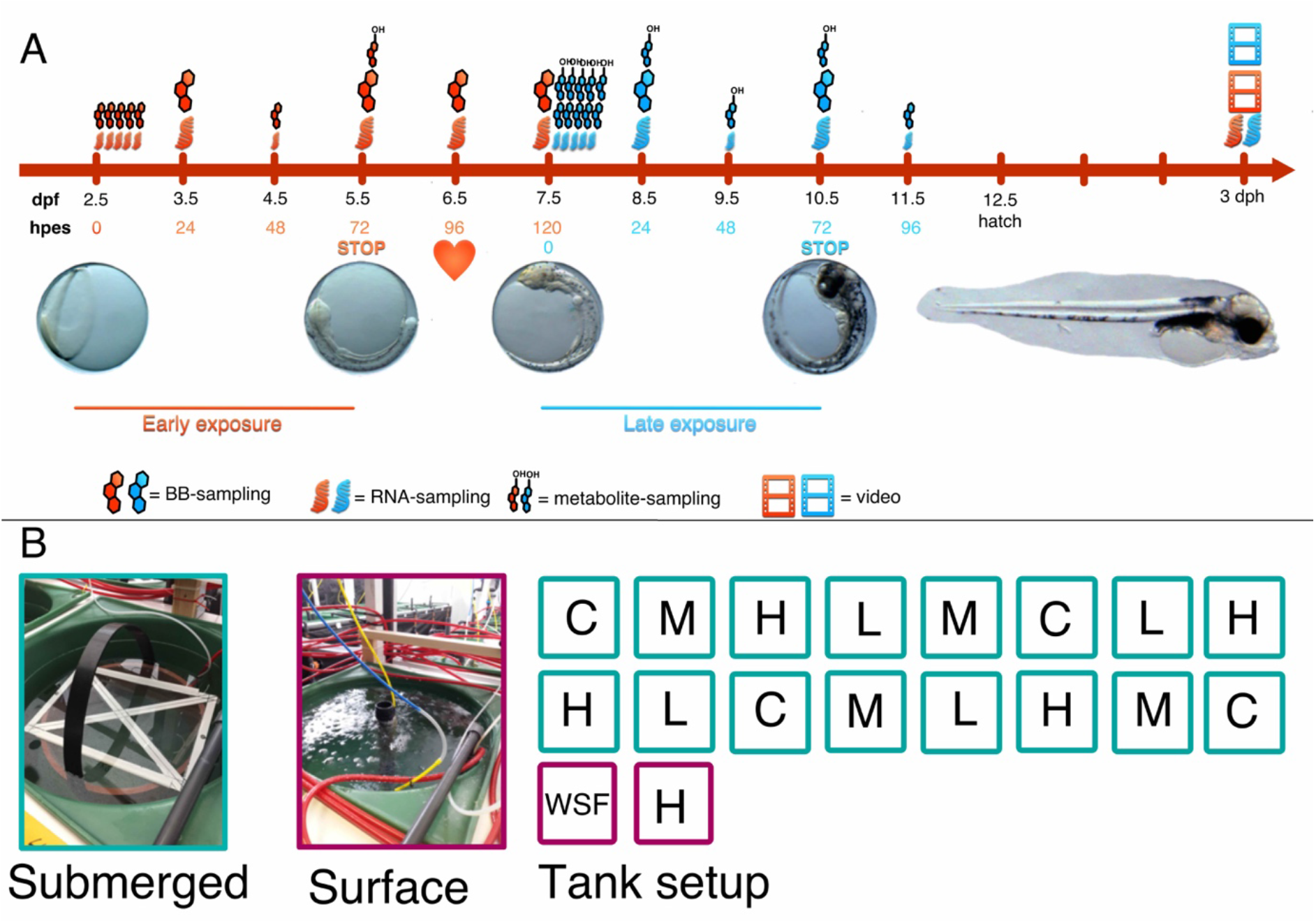
Experimental design and overview. A) Timeline for experiments in relation to development. Atlantic haddock (Melanogrammus aeglefinus) embryos were exposed from 2.5 days post fertilization (dpf) to 5.5 dpf (early exposure) and from 7.5 dpf to 10.5 dpf (late exposure). Heart symbol indicates time point for observation of first heartbeat (6.5 dpf). Samplings were performed from 1-120 hours post exposure start (hpes) for the early exposure, and from 1-96 hpes for the late exposure. Videos were conducted 3 days post hatching (dph) for both experiments. Red and blue labeling of timepoints and sampling symbols represent early and late exposure, respectively. Small and large sampling symbols indicate sampling of high doses and control and sampling of all groups, respectively. B) Tank setup. C = control, L = Low dose, M = Medium dose, H = High dose, WSF = Filtered oil dispersion (water soluble fraction). The turquoise tanks were submerged.

After exposure, ≈1000 eggs were transferred to recovery flow through tanks with clean water where they were kept until termination of experiment at 3 days post-hatching (dph). Throughout this work, multiple terms are used to define timepoints, based on days post-fertilization (dpf), hours post-exposure start (hpes), and days post-hatching (dph) (Fig 1A).

### 2.2. Sampling regime

Duplicate sample collection from all 18 tanks for total RNA extraction, body burden and metabolites occurred at 24, 72 hpes and 3 dph for both experiments, as well as 96 and 120 hpes for the *early* exposure. A more thorough time series with triplicate samples was collected from a tank of each of four selected treatments (*C, H sub, H surf* and *H WSF*) following the sampling regime described in Fig. 1A, with sampling points at 1, 2, 4, 8, 12, 24, 48, 72, 96 and 120 hpes. Fewer replicates were taken for *H surf* and *H WSF* at some of the latest time points due to high mortality. The precise number of eggs in each sample pool was counted from microscope pictures.

### 2.3. Analytical chemistry

#### 2.3.1. PAH analysis in exposure water and body burden samples

Water samples (1 L) were taken from each exposure tank at the beginning and the end of each experiment, preserved by acidification (HCl, pH < 2) and then stored at 4 °C in the dark until further handling. Details are given in Sørensen et al. (2017). In short, the water samples were extracted twice by partitioning to dichloromethane (30 mL) in a separatory funnel (2 min). Deuterated internal standards (naphthalene-d8, biphenyl-d8, acenaphtylene-d8, anthracene-d10, pyrene-d10, perylene-d12 and indeno[1,2,3-c,d]perylene-d12) were added prior to extraction to account for analytes loss during extraction. The combined extracts were concentrated by solvent evaporation prior to analysis by GC-MS. The following PAH compounds were measured: C0-, C1-, C2-, C3-, C4-naphthalenes (NAP), anthracene (ANT), acenaphthylene (AC), acenaphtene (AE), C0-, C1-, C2-, C3-fluorene (FLU), C0-, C1-, C2-, C3, C4-phenanthrene/anthracenes (PHE), C0-, C1-, C2-, C3-, C4-dibenzothiophenes (DBT), fluoranthene (FL), C0-, C1-, C2-, C3-pyrene (PYR), benzo[a]anthracene (BAA), C0-, C1-, C2-, C3, C4-chrysene (CHR), benzo[b+j+k]fluoranthene (BKF), benzo[e]pyrene (BEP), benzo[a]pyrene (BAP), perylene (PER), benzo[g,h,i]perylene (BP), indeno[1,2,3-c,d]pyrene (IND) and dibenzo[a,h]anthracene (DBA).

Samples for PAH body burden were collected frequently (Fig. 1A) to detect immediate changes in PAH uptake. The 48 hpes time point for the *late* exposure was compromised during analysis and data are not available. Each sample collected consisted of 50-100 pooled animals preserved by flash-freezing in liquid nitrogen and stored at −80 °C. Tissue samples were extracted by solid-liquid extraction and cleaned-up by solid phase extraction (SPE) prior to analysis. The method is described in detail in Sørensen et al. (2017). Briefly, after addition of internal standards the samples were homogenized in 2 mL dichloromethane-hexane (1:1, v/v), followed by addition of sodium sulphate (150 mg) and two rounds of vortex extraction (30 sec) and centrifugation (2000 rpm, 2 min). The combined organic extract was concentrated to ~1 mL prior to clean-up by SPE (Agilent Bond Elut SI, 500 mg). The SPE columns were conditioned with hexane and the extract eluted with 6 mL 10% dichloromethane in hexane. The purified extract was concentrated to 100 μL under a gentle stream of N_2_.

Water and body burden samples were analyzed on an Agilent 7890 gas chromatograph coupled to an Agilent 7010c triple quadrupole mass detector (GC-MS/MS) as described previously in Sørensen et al. (2017). For the body burden samples, a matrix-matched calibration was used, made from the blank extraction of haddock and cod eggs.

#### 2.3.2. Phenatharene metabolite analysis in haddock eggs

There are today no analytical methods that are able to detect the whole metabolome of PAH in fish (Beyer et al., 2010). In the present study we have included detection of phenathrene metabolites as representatives of complex PAH metabolism. We analyzed the main products of phenathrene metabolism in fish which are dominated by dihydroxylated-phenanthrene together with minor levels of monohydroxylkated-phenanthrenes (Goksoyr et al., 1987; Pampanin et al., 2016; Pangrekar et al., 2003; Sette et al., 2013).

The following standards of target compounds were purchased from Chiron AS (Trondheim, Norway): 1-hydroxyphenanthrene (1-OH-PHE), 2-hydroxyphenanthrene (2-OH-PHE), 3-hydroxyphenanthrene (3-OH-PHE), 4-hydroxyphenanthrene (4-OH-PHE), 9-hydroxyphenanthrene (9-OH-PHE), *trans*-9,10-dihydroxy-9,10-dihydrophenanthrene (9,10-diol-PHE) and surrogate standard 2-hydroxynaphthalene-d7 (2-OH-NPH-d7). The standard *trans*-1,2-dihydroxy-1,2-dihydrophenanthrene (1,2-diol-PHE) was purchased from MRI Global Carcinogen Repository (Kansas City, MO, USA). Samples for metabolite analysis were collected at 72 hpes in *early* exposure and 1-72 hpes in *late* exposure (Fig. 1A), flash frozen and kept at −80 °C until analysis. Between 50-100 eggs pooled from the same treatment were homogenized and a 50 μL aliquot of the homogenized mixture was diluted with water to 200 μL and spiked with surrogate standard. The homogenate was then added to phospholipid removal cartridges (Phenomenex Phree cartridges) contaning methanol and eluted under vacuum to remove phospholipids and proteins, followed by enzymatic hydrolysis by adding a pH-5 buffer containing 2000 U of β-glucuronidase/sulfatase (Sigma-Aldrich) and incubating for 1 hour at 40 °C. The hydrolysis reaction was then suspended with the addition of glacial acetic acid. We used SPE with Strata-X 200 mg/6 mL cartridges (Phenomenex Inc.) preconditioned with methanol and distilled water. Sample was added and washed with water and 63:37 methanol:water (v/v) before drying 30 minutes under vacuum. Target metabolites were finally eluted using 65:35 methanol:diethyl ether (v/v). Final volume was reduced to 3 mL before addition of internal standards and an aliquot was transferred for final analysis of the PAH metabolites.

Instrumental analysis was performed with a Waters Acquity high performance liquid chromatograph (LC) coupled with an AB Sciex QTRAP 5500 tandem mass spectrometer (MS) operating in negative electrospray ionization mode with ion spray voltage of 4500 V and multiple reaction mode (MRM) with the source temperature at 650 °C. The LC was equipped with a 0.2-μm pre-filter followed by a 2.1 x 5.0 mm (1.7-μm particle size) reversed phase guard column (BEH Shield RP18, Waters Co) and a 2.1 x 150 mm (1.7 μm particle size) reversed-phase column (BEH Shield RP18, Waters Co). Water (solvent A) and methanol (solvent B) were used as a mobile-phase with a 29-minute linear gradient as follows (solvent A/solvent B): initial gradient was 50/50 at 0.2 mL/min; 15 min to 20/80 at 0.2 mL/min; 0.1 min to 100% solvent B and 0.1 min to 0.3mL/min; held at 100% B for 6.8 min at 0.3 mL/min; 0.5 min to return to initial gradient 50/50 at 0.3 mL/min; held at 50/50 for 6.0 min at 0.3 mL/min; 0.1 min to 0.2 mL/min and kept for 0.4 min. The column temperature was maintained at 45°C.

Sum OHPHEs (ΣOHPHEs) is the sum concentrations of 1-OH-PHE, 2-OH-PHE + 3-OH-PHE (estimate concentration, as these are chromatographically co-eluting isomers), 4-OH-PHE and 9-OH-PHE. Sum PHE-diols (ΣOHPHEs) is the sum concentrations of 1,2-diol-PHE and 9,10-diol-PHE. Sum PHE metabolites (Σ PHE metabolites) is the sum concentrations of all metabolites measured. The quantitative results were converted from ng metabolites/embryo to ng/g by using average mass of one haddock egg (0.00218 g).

### 2.4. Detection of Cyp1a activity

Ethoxyresorufin-O-deethylase (EROD) assay was applied to detect Cyp1a activity. Activity was measured at 72 hpes (5.5 dpf) in the *early* exposure. The embryos were at the end of Organogenesis 1, at the 10-30 somite/cardiac cone developmental stage (Fridgeirsson, 1978; Hall et al., 2004; Sørhus et al., 2016a). The procedure was performed as described in Gonzalez-Doncel et al. (2011). Briefly, 7-ethoxyresorufin (7ER) was added to live embryos and the Cyp1a-catalyzed fluorescing product, resorufin, was detected with fluorescence microscopy. A stock of 7ER (48 mg 7ER/L) (Sigma-Aldrich, Germany) was made in 100% DMSO (Thermo Fisher Scientific). Control, DMSO and crude oil exposed embryos were transferred to 24-well plates where they were incubated in the dark for 1 hour in 20 μg 7ER/L or DMSO in sterile sea water. After exposure the solution was drained off and exchanged with sterile sea water. Images of 12-20 embryos per treatment were captured with a digital camera (SPOT Insight 5 Mpx) coupled to a Nikon AZ100 fluorescence microscope. For each set of embryos, two fluorescence images were captured using the rhodamine filter (Ex 545/25 nm and Em 605/70 nm) to detect EROD activity and DAPI filter (Ex 350/50 nm and Em 460/50 nm) to detect the internal fluorescence of PAH compounds. In addition, a corresponding brightfield image was captured for each set of fluorescence. Exposure time and gain for all fluorescence images were set at 300 ms and 4 g. Semi-quantitative measurements of EROD fluorescence in the embryos was estimated using the adjusted light intensity in the whole egg in the various treatments. Adjusted light intensity was measuered using ImageJ software (Schneider et al., 2012; http://imagej.nih.gov/ij/, 1997–2016) and was calculated using the following formula: Adjusted light intensity = (light intensity in Treatment with 7ER)/(light intensity in Treatment with DMSO).

### 2.5. Imaging of live embryos/larvae and measurements of cardiac function

Images were taken at all sampling points to define developmental stage, morphological abnormalities and degree of oil droplet accumulation. At 3 dph, all groups were imaged to assess morphological and cardiac functional endpoints. Images were collected for all exposure tanks in both *early* and *late* exposures. Digital still micro-images and 20-second videos of live 3 dph larvae were obtained using an Olympus SZX-10 Stereo microscope equipped with a 1.2 megapixel resolution video camera (Unibrain Fire-I 785c) controlled by BTV Pro 5.4.1 software (www.bensoftware.com). Image calculations were calibrated with a stage micrometer.

Animals were immobilized in a glass petri dish filled with 3 % Methylcellulose (Sigma-Aldrich, Germany) in seawater and kept at 8°C using a temperature-controlled microscope stage (Brook Industries Inc. Lake Villa, IL. USA). Length of ethmoid plate, mandible, area of edema and length, systolic and diastolic diameter of ventricle and atrium were measured using ImageJ (Schneider et al., 2012; http://imagej.nih.gov/ij/, 1997–2016) with the ObjectJ plugin (https://sils.fnwi.uva.nl/bcb/objectj/index.html) from images as described previously (Sørhus et al., 2016b). Additionally, eye diameter was quantified, and eye shape and deformities were described according to categories: normal shape (no), bend shape (be), irregular shape (irr), protruding lens (pl). Six distinct craniofacial phenotypes were identified. Degree of looping of heart was also described (normal looping (n), poor looping (p) and severly poor looping (sp)). The area occupied by edema fluid was quantified as the difference between the total area of the yolk sac and the area of the yolk mass, expressed as a percentage of total yolk sac area. Similarly, ventricular and atrial diastolic (D) and systolic diameter (S) were used to estimate the fractional shortening (FS = (D – S)/D). Measurements from both images and videos were performed blind.

### 2.6. Oil induced gene expression changes

All animals collected for RNA extraction were frozen in liquid nitrogen and stored at −80 °C after being imaged (as described in the previous section). Total RNA was extracted from frozen pools with 10 eggs/larvae with Maxwell HT simplyRNA (Promega corporation) according to the manufacturer’s instructions using Biomek 4000 Automated liquid handler (Beckman coulter). Quantity and quality of the extracted total RNA was checked using Nanodrop spectrophotometer (NanoDrop Technologies, Wilmington, DE, USA) and Bioanalyzer (Agilent Technologies, Santa Clara, CA, USA) respectively. cDNA was subsequently generated using SuperScript VILO cDNA Synthesis Kit (Life Technologies Corporation), according to the manufacturer’s instructions, and was normalized to obtain a concentration of 50ng/μL. The described frequent sampling regime was followed to detect immediate gene expression changes (Fig. 1A and see the previous section “*Sampling regime*”).

Specific primers and probes for real-time qPCR analysis of Atlantic haddock were designed with Primer Express software (Applied Biosystems, Carlsbad, California, USA) (*cyp1a* and the technical reference *ef1a*) or Integrated DNA Technology (IDT) probe and primer design software (IDT Inc., Iowa, USA) (*cyp1b, cyp1c, cyp1d, abcb1, rhag* and technical reference *rxrba*), according to the manufacturer’s guidelines. Primer and probe sequences are given in Supplementary table S1. Real-time qPCR assays were performed in duplicate, using 384-well optical plates on a QuantStudio 5 Real Time PCR System (Thermofisher Scientific) with settings as follows: 95 °C for 20 s, followed by 40 cycles of 95°C for 1 s and 60°C for 20 s. Duplicates with standard deviation^2^ (SD^2^ ≥ 0.05 were either rerun or eliminated from the dataset. No-template control, reverse transcriptase enzyme control and genomic DNA controls were included. For each 5 μl PCR reaction, a 1.5 μl cDNA 1:40 dilution (1.9 ng) was mixed with 200 nM fluorogenic probe, 900 nM sense primer, 900 nM antisense primer in 1xTaqMan Fast Advanced Master Mix (Applied Biosystems, Carlsbad, California, USA). Gene expression data were calculated relative to control in the first sampling point using the ΔΔΔCt method generating reference residuals from *ef1a* and *rxrba*, as described in detail in Edmunds et al. (2014).

### 2.7. Localization of *rhag* gene expression

Location of *rhag* gene expression was examined using a method derived from previous protocols (Hall et al., 2003; Thisse and Thisse, 2008; Valen et al., 2016). We performed whole mount *in situ* hybridization on unexposed 10 dpf embryos to determine location of *rhag* in early life stages of fish. Detailed protocol can be found in Supplementary information.

### 2.8. Statistics

Statistical differences in Cyp1a activity, morphology, cardiac function, PAH water concentration, PAH body burden, and OHPAH metabolites were tested using one-way ANOVA with Dunnet’s multiple comparison in R (The R Foundation for Statistical Computing Platform). In the gene expression data, statistically significant differences were tested using both One-way and Two-way ANOVA with Dunnet’s multiple comparison in R. for? PAH water concentration, PAH body burden, OHPAH metabolites and gene expression data were log-transformed due to large differences in levels and thus standard deviation. In craniofacial and eye phenotypes and appearance of silent ventricle, statistical differences between treatments and control were evaluated using chi-square test. Statistical difference was denoted p<0.05 (*), p<0.01 (**), p<0.001 (***). Error bars in all figures represent the standard deviation of the mean.

### 2.9. Ethics statement

The animals were monitored daily, and any dead larvae were removed. All sampled embryos and yolk sac larvae were euthanized immediately in liquid nitrogen. The Austevoll Aquaculture Research station has the following permission for catch and maintenance of adult Atlantic haddock: H-AV 77, H-AV 78 and H-AV 79. These are permits given by the Norwegian Directorate of Fisheries. Furthermore, the Austevoll Aquaculture Research station has a permit to run as a Research Animal facility using fish (all developmental stages), with code 93 from the national IACUC; NARA, although no approval is necessary to perform studies with fish embryos and yolk sac larvae.

## 3. Results

Overall, both *early* and *late* exposure resulted in functional and morphological abnormalities, but they were more severe in the *early* exposure. Tissue uptake of PAHs was higher in the *early* exposure, followed by a stronger, but delayed *cyp1a* induction. The *H WSF* group showed very few oil induced abnormalities in both exposures. Gene expression patterns were unique between *early* vs *late* exposures.

### 3.1. Oil droplet fouling in surface and submerged embryos

Oil droplet fouling was observed on embryos in both submerged (*L, M, H sub*) and surface (*H surf*) exposures regardless of timing, and was evident after only 1 hour of exposure (Fig 2A and B, top panels). The oil droplet fouling effect was higher in the surface (*H surf*) exposure due to the accumulation of oil microdroplets on the surface (Fig. 2). While oil droplets in the *early* exposure were distributed all over the surface, the *late* exposure embryos mainly accumulated oil droplets at one area on the chorion (Fig 2B, filled black arrows). In the filtered high dose group (*H WSF*), no oil droplets were detected on the chorion (Fig 2A and B, right panel). After transfer to clean water, some of the droplets were washed off, however, in addition to a reduced number of apparent oil droplets (brown), translucent droplets were now visible on the surface (white arrow head Supplementary Fig. S1). This phenomenon was not observed in the *late* exposure due to hatching that occurred shortly after end of exposure (11 dpf).

**Figure 2:**
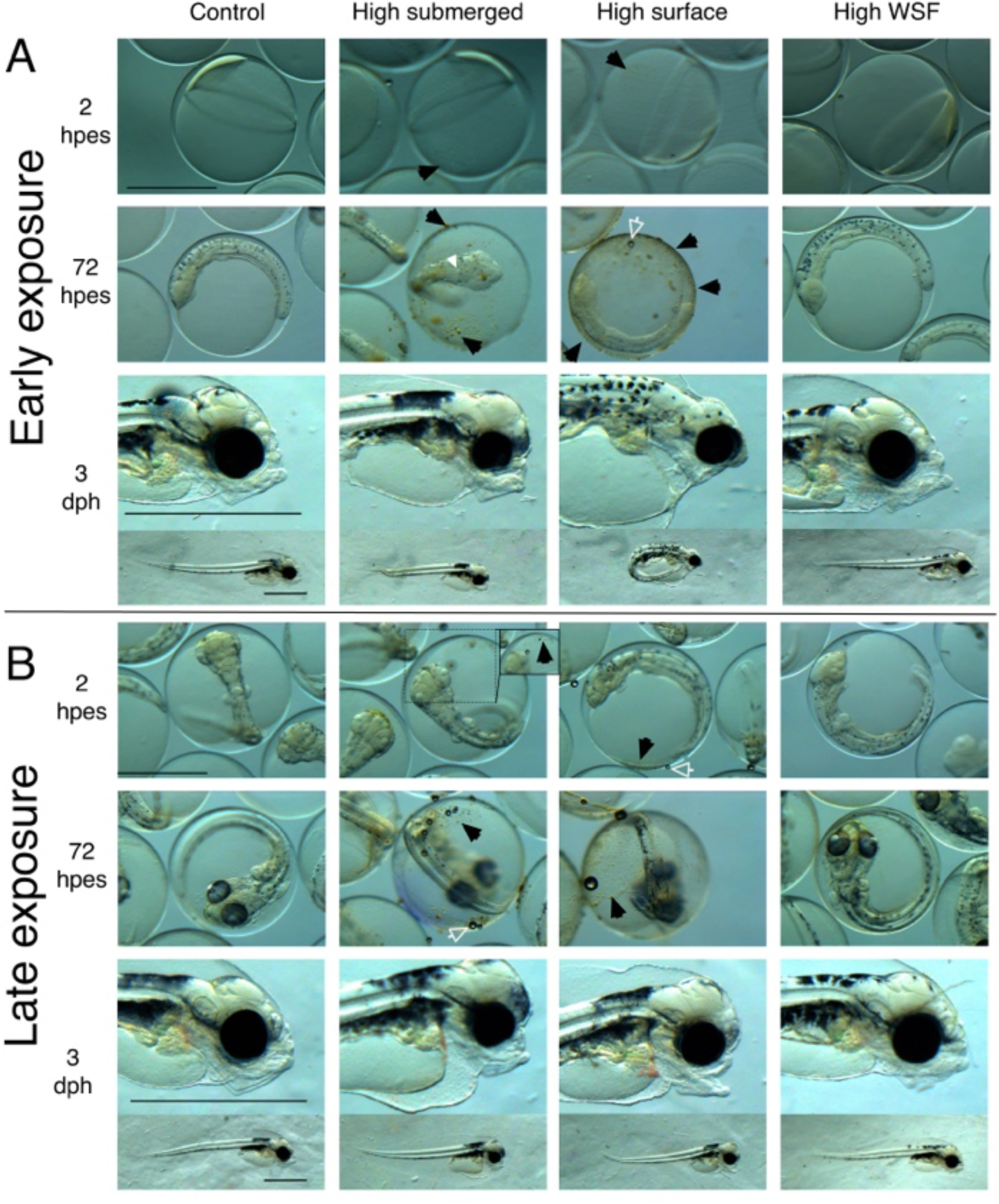
Oil droplet fouling and morphology. A) early and B) late embryonic exposure of Atlantic haddock. Top panels show 2 hours post exposure start (hpes), middle panels 72 hpes (exposure stop) and lower panels display representative phenotypes at 3 days post hatching (dph). Black arrows: oil droplets. White arrow head: indicates cardiac cone. White open arrow: air bubble trapped on the chorion. WSF: water soluble fraction. Scale bar: 1000 μm.

### 3.2. Oil exposure and PAH uptake

#### 3.2.1. PAH water and tissue concentration

We successfully delivered doses of dispersed crude oil to the exposure tanks during the experiments. Total PAHs (dissolved and droplet associated) were monitored at the beginning and end of each exposure experiment (Fig. S2). The water concentration of PAH was lower in the *H WSF* than the other high dose groups, and reflected the PAH contribution from the oil droplets. There were some differences in the total PAH concentration between the two experiments, but there were a clear gradients among the different treatment in both studies: total PAH concentration (μg PAH/L): Average (start and stop) concentrations after *Early* exposure *C* = 0.04 +/- 0.01, *L* = 0.47 +/- 0.61, *M* = 2.3 +/- 2.4, *H sub* = 5.2 +/- 1.7, *H surf* = 6.4 +/- 0.8, *H WSF* = 3.7 +/- 1.0. *Late* exposure C = 0.04+/-0.01, L = 0.5+/-0.2, M = 2.2+/-0.8, *H sub* = 9.7 +/- 2.7 *H surf* = 8.3 +/- 2.9, *H WSF* = 6.3 +/-0.9

An immediate and prominent uptake of PAHs into the embryo was detected (Fig. 3). Body burden measurements of all doses at all time-points in both experiments had statistically significant elevated levels of PAH compared to control. Average total PAH tissue concentrations in the exposed groups for *early* exposure ranged from 38 (1 hpes, *H WSF*) to as high as 7800 ng/g ww (72 hpes, *H surf*). Values in the *late* exposure were between 75 (24 hpes, *L*) and 3000 ng/g ww (72 hpes, *H surf*). Notably, despite higher water concentrations in the *late* exposure, the highest tissue concentrations were found in the *early* exposure. Moreover, the tissue concentration in the *early* exposure peaked at a later time point than in the *late* exposure (72 hpes vs 24 hpes). The levels in the *H WSF* groups (in which the water was filtered) were more similar to the medium group, despite sharing the same oil content of the high dose (300 μg oil/L) (Fig. 3A). Figure 3B shows the profile of PAHs at 72 hpes, where, notably, the high molecular weight (HMW) PAH group (inset in Fig. 3B) found in medium, *H sub* and *H surf* groups, was lacking in the *H WSF* group. These HMW PAHs, *e.g*. PYRs and CHRs, accumulated substantially more in the *H surf* than in other groups. In addition, in the *H surf*, the levels of these HMW PAHs were up to four times higher in *early* vs *late* exposure (Fig. S3). Detailed profiles for individual PAHs at each time point are documented in Supplementary Dataset S1.

**Figure 3:**
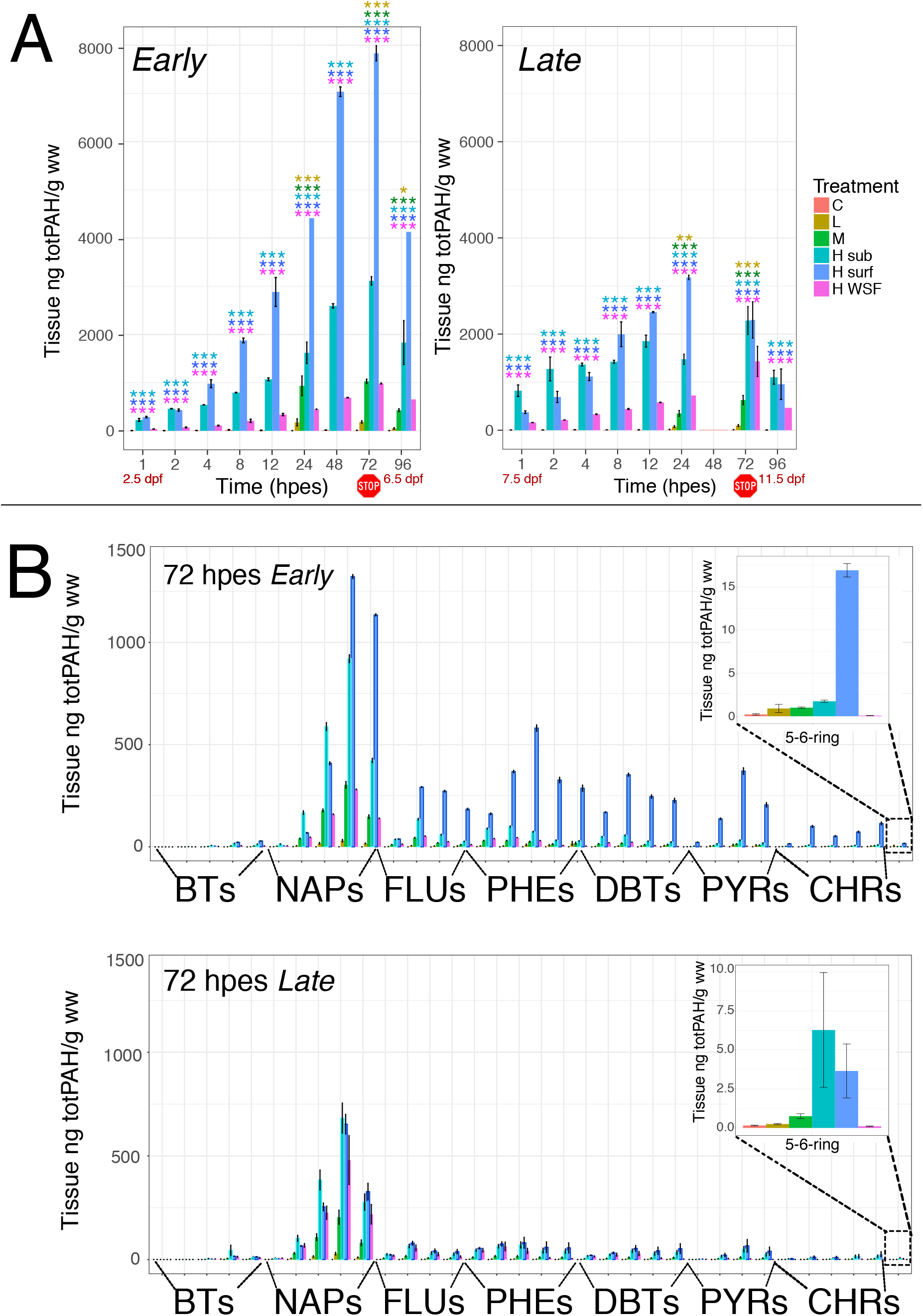
Tissue uptake of polycyclic aromatic hydrocarbons (PAHs). A) Total body burden of PAHs from 1-96 hours post exposure start (hpes) in early (Left panel) and late (right panel) exposure. Exposure stop was after 72 hours. Statistical difference from control was tested at each time point with One-way ANOVA and significant differences are indicated by **=p<0.01 or ***: p<0.001. Note, at 96 hpes only one replicate from H surf and H WSF (early exposure) and H WSF (late exposure) were analyzed. B) PAH profile at exposure stop (72 hpes) in early (left panel) and late (right panel) exposure. Inset: magnification of 5-6 ring distribution in the treatment groups. Dpf: days post exposure, C: control, L: low dose, M: medium dose, H sub: high dose submerged, H surf: high dose surface, H WSF: high dose water soluble fraction, BTs: Benzothiophenes, NAPs: Naphtalenes, FLUs: Fluorenes, PHEs: Phenanthrenes, DBTs: Dibenzothiophenes, PYRs: Pyrenes, CHRs: Chrysenes.

#### 3.2.2. Hydroxylated phenanthrene metabolites

Hydroxylated phenanthrene metabolites were analyzed in the control and three high dose groups (*H sub*, *H surf* and *H WSF*) at *early* (only at exposure stop, 72 hpes) and *late* exposure (1-72 hpes). Metabolites assessed are presented as sum of monohydroxyphenanthrenes (ΣOHPHE) and sum of dihydrodiol phenanthrenes (ΣPHE-diols) (Fig. 4). Within these groups 1,2-diol-PHE was the dominating metabolite and accounted for approximately 90% of total detected metabolites (average of all treatments). Of the ΣOHPHE, 1-OH-PHE was the most abundant and contributed with 9% of the total metabolites (Supplementary Dataset S2).

**Figure 4:**
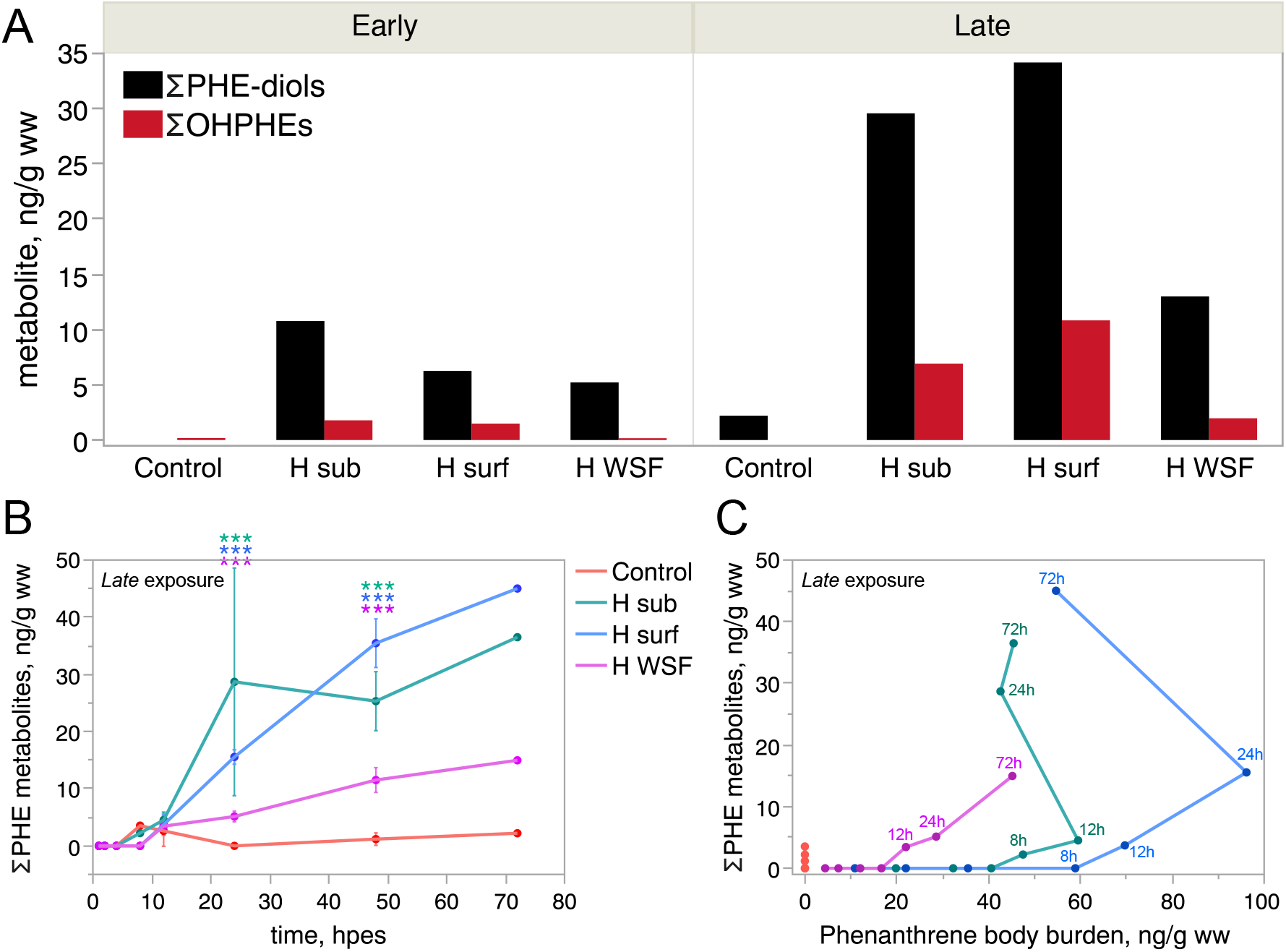
Tissue concentration of phenanthrene metabolites. A) Concentrations of PHE-diols and OHPHEs at 72 hpes in early and late exposures (one sample per treatment). B) Concentrations of ΣPHE metabolites over time in late exposure. Statistical difference from control was tested at 12 hpes, 24, hpes and 48 hpes with one-way ANOVA with Dunnet’s multiple comparison, and significant differences are indicated by ***: p<0.001. C) Sum of phenanthrene metabolites versus mean body burden of phenanthrene in late exposure. The backwards curve suggests that metabolism is faster than uptake towards the end of the exposure. H sub: high dose submerged, H surf: high dose surface, H WSF: high dose WSF.

Metabolite levels at exposure stop (72 hpes) for *early* exposure were approximately 3 times lower than *late* exposure (Fig 4A). In the *late* exposure, where we analysed for a metabolite profile over time, we observed significant levels of phe**n**anthrene metabolites in all high dose groups after 24 hpes, and these levels continued to increase until exposure stop (Fig. 4B). The maximum body burden of PHE metabolites was found in the *H surf* group (72 hpes) in a concentration of 45 56ng/g ww, which is in the similar concentration ranges as the body burden of the parent PHE (95 ng/g ww found after 24 hpes). The metabolites in the *H WSF* were approximately 1/3 that of *H sub*, although they followed a similar trend over time. (Fig. 4B). The correlation between phenanthrene metabolites and body burden (Fig. 4C) demonstrates that the rate of biotransformation exceeds the rate of PAH uptake. By 12 hpes (*H sub*) and 24 hpes (*H surf*), the curve turns back on itself indicating that the formation of metabolites is increasing while the PAH body burden has begun to decrease (even under continuous exposure).

### 3.3. Temporal expression of *cyp1a, cyp1b, cyp1c* and *cyp1d* and EROD activity

#### 3.3.1. Temporal expression of Cyp1s

Expression of *cyp1a* was immediate in both experiments. It was increasing throughout the *early* exposure (Fig. 5A),and the expression stayed steady from 12-72 hpes at the late exposure (Fig 5B). In the *early* exposure a 5- and 6-fold significant change of *cyp1a* was detected after 8 hours of exposure in *H sub* and *H surf*, respectively (Fig. 5A). All treatments at investigated time-points were up-regulated from 12 hpes (Fig. 5A). As with PAH body burden, the *cyp1a* expression in the *H WSF* group fell between the levels detected in low and medium dose in all time points except at 120 hpes, which reflects the toxicity contribution of oil droplets in the exposure that is reduced when the exposure water is filtered to yield the WSF exposure. In the *late* exposure, the induction of *cyp1a* was much more rapid with 2 fold up-regulation after only 2 hours of exposure in the *H surf* group (Fig. 5B). From 8 hpes and onward significant up-regulation of *cyp1a* was seen in all treatment groups including low and medium at 24 and 72 hpes (Fig. 5B). Similar to the *early* exposure, the *H WSF* group *cyp1a* expression fell between levels detected in the medium and low dose groups (Fig. 5B).

**Figure 5:**
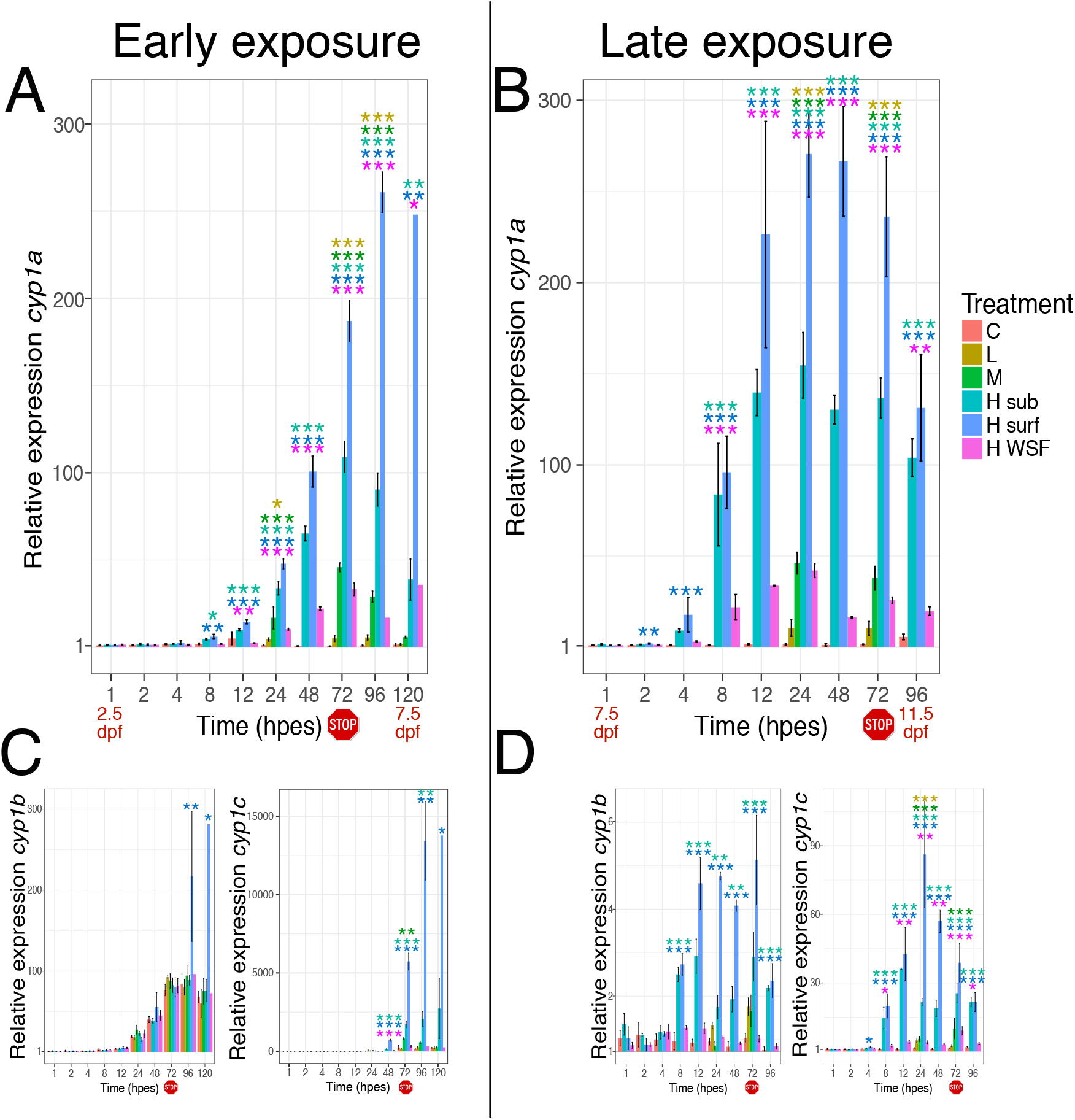
Relative expression of cyp1s. Relative expression of cyp1a after early (A) and late (B) exposure. C) shows relative expression of cyp1b (left panel) and cyp1c (right panel) after early exposure, while D) displays cyp1b (Left panel) and cyp1c (right panel) after Late exposure. Expression was related to control group at exposure start to detect both developmental and treatment related expression. Exposure stop is indicated with a stop icon. Statistical difference from control at each time point was tested with One-way ANOVA with Dunnet’s multiple comparsions and significant differences are indicated by *=p<0.05, **=p<0.01 or ***: p<0.001. At 120 hpes in the early exposure, only one sample was collected for H surf and H WSF. Hpes: hours post exposure start, Dpf: days post exposure, C: control, L: low dose, M: medium dose, H sub: high dose submerged, H surf: high dose surface, H WSF: high dose WSF.

Induction of *cyp1a* correlated with tissue uptake of PAH (Fig. S4). However, one of the low dose tanks, L4, showed low PAH uptake and *cyp1a* induction in both *early* and *late* exposures. It was therefore considered an outlier and eliminated for further analysis in both experiments. Embryos from the L4 tank clustered to the control tanks rather than the low dose tanks at both 24 hpes and 72 hpes (Fig. S5, 72 hpes).

The expression of the other *cyp1* paralogs, *cyp1b* and *cyp1c*, was delayed in the *early* exposure and genereally followed the same pattern as *cyp1a* expression in the *late* exposure (Fig. 5). The expression profile over time for the control group show a highly dynamic temporal expression of *cyp1b, cyp1c* and *cyp1d* while *cyp1a* showed a stable expression throughout embryonic development (Fig. S6A-D). In the early exposure, *cyp1c* was not appreciably expressed regardless of treatment until 48 hpes (Fig. 5C) and at 72 hpes a developmentally related 200 fold up-regulation in the control group was observed (Fig. S6). Similarly, a general up-regulation of *cyp1b* was seen from 24 hpes, with a significant upregulation in surface dose at the two latest time points (Fig. 5C). In comparison, at peak at 96 hpes *H surf* showed a 221 fold up-regulation of *cyp1a* while *cyp1b* and *cyp1c* showed a 2.5 fold and 82 fold up-regulation, respectively, compared to control at 96 hpes. For *cyp1d*, almost no transcripts were detected at any of the time points in either dose/control in the *early* exposure. In the *late* exposure *cyp1d* was generally up-regulated at 72 hpes, with a significant up-regulation in *M* and *H sub* groups (Fig. S7). After 10 days of recovery (*early* exposure) at 3 dph one of the groups *H surf*, still had significant up-regulation of one of the *cyp1s*, namely *cyp1b* (Fig. S8A-D, left panel). While at 3 dph after 5 days of recovery (*late* exposure), significant up-regulation in *H surf* group was observed for all *cyp1s*, and in *H sub* group for *cyp1a, b* and *c* (Fig. S8A-D, right panels).

#### 3.3.2. Cyp1a activity

Ethoxyresorufin-O-deethylase (EROD) assay was applied to detect Cyp1a activity and was measured at 72 hpes (5.5 dpf) in the *early* exposure. Cyp1a activity was detected in the skin, mainly in the trunk, and in the otoliths in the *H sub* and *H surf* exposures. Low and medium doses showed EROD activity in the trunk, while very little activity was detected in *H WSF* (Fig. S9). No specific fluorescence was detected in the embryos incubated in DMSO (images not shown). The internal fluorescence activity of the PAHs in the oil droplets are shown in Figure S9C.

### 3.4. Crude oil induced malformations

Craniofacial, eye and growth abnormalities were seen in both exposures (Table 1). We defined six distinct dose-dependent craniofacial phenotypes, given here in increasing severity: mild Bulldog (MBD) and Bulldog (BD) with reduced upper jaw, Upper Jawbreaker (UJB) with a posteriorly twisted upper jaw, Total Jawbreaker (TJB) with posteriorly twisted jaws in general, Darth Vader (DV) with an underdeveloped upper jaw and hanging lower jaw and Hunchback (HB) with severe reduction of all jaw structures with reduced brain structures, abnormal finfold, spinal curvature and poor dorsal rotation of the head (Fig. 6).

**Table 1:**
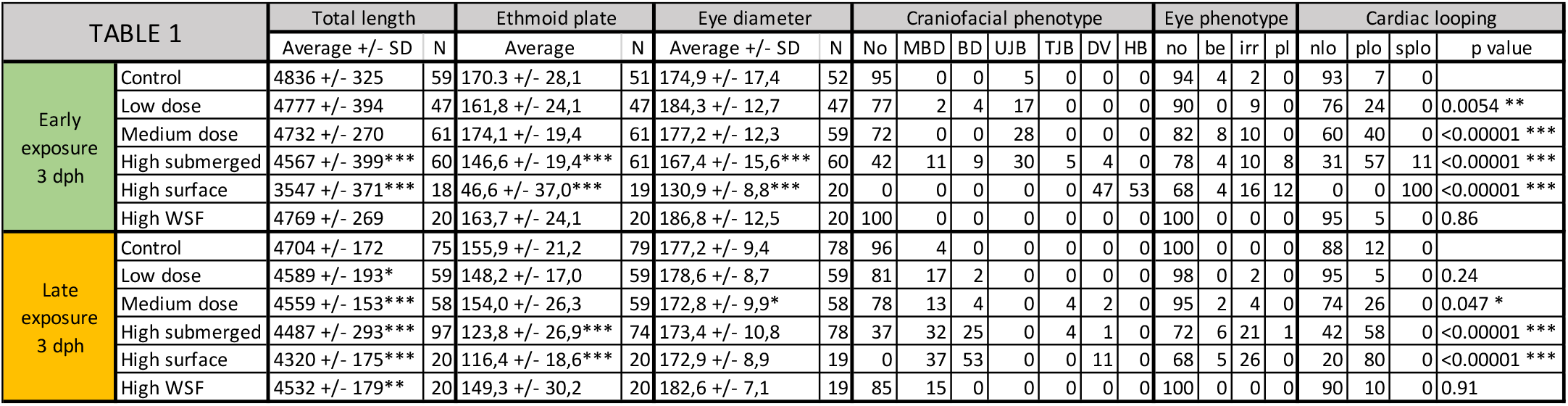
Morphological abnormalities after early and late embryonic exposure. WSF; water soluble fraction, SD; standard deviation, N; total number of animals analyzed, No; normal phenotype, MBD; mild bulldog phenotype, BD; bulldog phenotype, UJB; upper jawbreaker phenotype, TJB; total jawbreaker phenotype, DV; Darth Vader phenotype, HB; hunchback phenotype, no; normal eye shape be; bend eye shape, irr; irregular eye shape, pl; protruding lens, nlo; normal looping, plo; poor looping, splo; severely poor looping. Statistical significant difference from control is indicated by asterisks; p<0.05 *, p<0.001 **. P<0.0001 ***.

**Figure 6:**
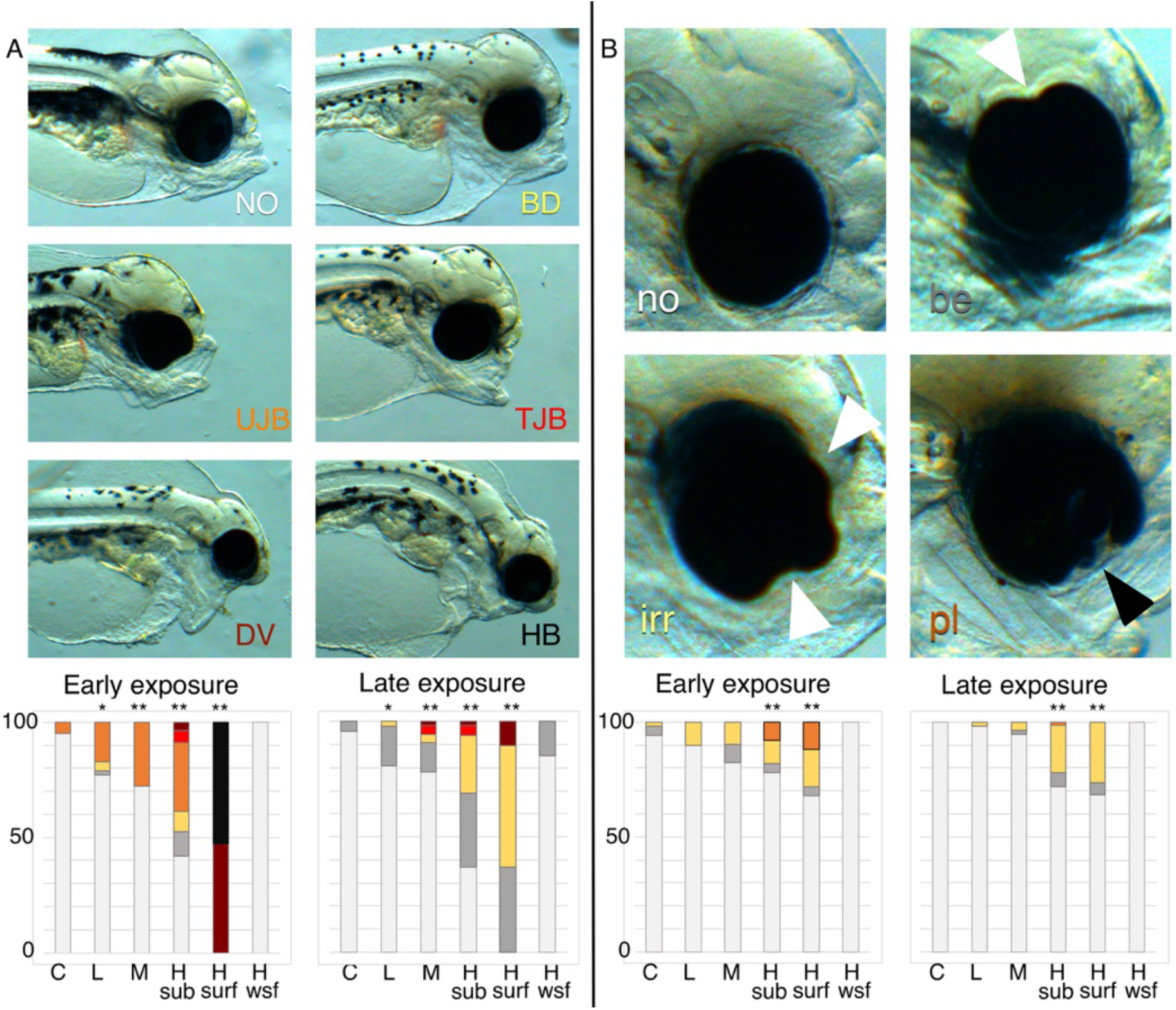
Oil induced craniofacial and eye phenotypes. A) Oil induced craniofacial phenotypes, NO: normal, BD: Bulldog, UJB: Upper jawbreaker, TJB: Total jawbreaker, DV: Darth Vader, HB: Hunchback. The bottom panel shows the distribution of the craniofacial phenotypes after early and late exposure at 3 days post hatching. Coloration code: white: NO, grey: mild BD, yellow: BD, orange: UJB, red: TJB, dark red: DV, black: HB. B) Oil induced eye phenotypes, no: normal eye shape (white), be: bend shape (grey), irr: irregular shape (yellow), pl: protruding lens (orange). Bottom panel shows the distribution of eye phenotypes in early and late exposures. White arrow heads indicate irregularities in the eye shape, and black arrow head indicate protruding lens. Statistical difference from control was tested with Chi-square, and p<0.01 is indicated by **. C: control, L: low dose, M: medium dose, H sub: high dose submerged, H surf: high dose surface, H WSF: high dose water soluble fraction.

In the *early* exposure, most severe craniofacial malformations were observed in the *H surf* group (Fig. 6A, lower panel Table 1), where all animals were characterized as DVs or HBs (Fig. 6A). The *H sub* group possessed fewer malformations (50%) and the group comprised of BD/MBD, UJB and TJB animals and a few DVs (Fig. 6A, Table 1). In the *H WSF* dose, no malformed phenotypes were observed while low and medium dose both had occasions of the milder phenotypes (MBD/BD and UJB, 28%) (Fig. 6A, Table 1). The abnormal jaw phenotypes were linked to the shorter ethmoid plate. In accordance with the phenotypic observations, both *H sub* and *H surf* groups with the most severe jaw effects also showed a significantly reduced ethmoid plate (14% and 72% shorter, respectively). While the *H WSF*, low dose and medium dose showed no significant reduction in ethmoid plate compared to control (Table 1). Similarly, the total length of the larvae were only significantly reduced in the *H sub* and *H surf* groups (Table 1).

In the *late* exposure, the craniofacial malformations were generally milder, and the difference between the *H sub* and *H surf* groups were less evident (Fig. 6). The same phenotypes were present (MBD/BD, TJB and DV) but with an increased proportion in the surface group where all had craniofacial malformations (Fig. 6, Table 1). The *H WSF* group had approximately the same occurrence of the mildest phenotype (MBD/BD, 15 vs 19%) as the low dose (Table 1). In comparison, the medium dose had slightly lower occurrence of the mildest phenotypes (MBD/BD, 17%), but additionally had incidents of the more severe phenotypes (TJB and DV, 6%) (Table 1). Along with craniofacial phenotype, the ethmoid plate was significantly reduced in the *H sub* and *H surf* groups (Table 1). Total length of the larvae was reduced in all exposed groups (Table 1).

Eye shape and diameter were affected by oil exposure at both time points. Four eye phenotypes were observed: no phenotype, normal round shape (no), bend shape (be), irregular shape (irr) and protruding lens (pl) (Table 1). The frequency of any eye phenotype was higher in the *early* exposed animals, and the most severe phenotype, protruding lens, was only found in *H sub* (8%) and *H surf* (12%) groups. No effect on eye shape was seen in *H WSF* regardless of timing of exposure (Table 1). Severe reduction in eye diameter was found in the *early* exposed *H surf* and *H sub* animals (Table 1). Less effect on eye size was observed in the *late* exposure, only medium showed significantly reduced eye diameter.

### 3.5. Cardiac malformations

#### 3.5.1. Morphological abnormalities

The size of the heart was reduced in both exposures, but only *early* exposure resulted in severly malformed hearts with poor looping (Table 2). After *early* exposure, an effect on atrial dimensions was only detected in the *H surf* and *H sub* group (Table 2). The ventricular diastolic diameter was reduced in all exposure groups (9-43%) except for *H WSF*, and the *H surf* group also showed reduced ventricular systolic diameter (38%) (Table 2). There was no effect on atrial dimensions after *late* exposure in any of the doses. However, ventricular systolic and diastolic diameters were reduced in both *H sub* (12% and 19%, respectively) and *H surf* groups (19% and 31%, respectively) in the *late* exposure (Table 2). The hearts were not as malformed as in the *early* exposure, and no incidences of severly poor looping was observed (Table 2).

**Table 2:** Cardiac abnormalities after early and late embryonic exposure. WSF; water soluble fraction, AS; atrial systolic diameter, N; Number of animals analyzed, AD; Atrial diastolic diameter, VS; Ventricular systolic diameter, VD; Ventricular diastolic diameter, AFS; Atrial fractional shortening, VFS; ventricular fractional shortening, BPM; Beats per minute, SiV; Silent ventricle. Statistical significant difference from control is indicated by asterisks; p<0.05 *, p<0.001 **. P<0.0001 ***.

| TABLE 2 |  | AS |  | AD |  | VS |  | VD |  | AFS % |  | VFS % |  | Edema |  | Yolk sac |  | BPM |  | SiV % |  |
| --- | --- | --- | --- | --- | --- | --- | --- | --- | --- | --- | --- | --- | --- | --- | --- | --- | --- | --- | --- | --- | --- |
|  |  | Average +/- SD | N | Average +/- SD | N | Average +/- SD | N | Average +/- SD | N | Average +/- SD | N | Average +/- SD | N | Average +/- SD | N | Average +/- SD | N | Average +/- SD | N | % | N |
| Early exposure 3 dph | Control | 101.4 +/- 14.3 | 42 | 121.7 +/- 18.4 | 42 | 89.9 +/- 13.8 | 54 | 105.8 +/- 19.7 | 54 | 17.8 +/- 6.9 | 42 | 14.4 +/- 6.9 | 54 | 15 +/- 8.3 | 61 | 21.3 +/- 8.1 | 61 | 93.1 +/- 8.7 | 61 | 2 | 54 |
|  | Low dose | 94.0 +/- 15.3 | 20 | 115.4 +/- 18.4 | 20 | 85.9 +/- 9.1 | 34 | 96.0 +/- 13.3* | 34 | 18.1 +/- 8.2 | 20 | 9.8 +/- 6.9* | 32 | 12.3 +/- 6.1 | 48 | 24.3 +/- 9.8 | 48 | 90.4 +/- 7.4 | 38 | 13** | 48 |
|  | Medium dose | 93.5 +/- 13.9 | 22 | 113.9 +/- 17.1 | 22 | 85.4 +/- 13.5 | 50 | 94.1 +/- 15.2** | 50 | 20.2 +/- 8.8 | 22 | 8.8 +/- 7.8** | 50 | 13.3 +/- 8.9 | 61 | 23.7 +/- 10.7 | 61 | 89.2 +/- 9.5 | 53 | 24*** | 51 |
|  | High submerged | 91.5 +/- 18.2* | 42 | 114.6 +/- 20.2*** | 44 | 84.1 +/- 15.6 | 57 | 91.7 +/- 19.5*** | 57 | 19.8 +/- 8.9 | 42 | 7.8 +/- 9.1*** | 57 | 17.6 +/- 10.9 | 61 | 24.1 +/- 11.2 | 61 | 90.5 +/- 10.3 | 64 | 20*** | 40 |
|  | High surface | 86.2 +/- 14.0** | 17 | 101.0 +/- 18.6 | 17 | 56.1 +/- 20.1*** | 15 | 59.8 +/- 20.1*** | 15 | 13.9 +/- 9.0 | 17 | 5.5 +/- 12.4*** | 15 | 30.0 +/- 13.1*** | 18 | 42.5 +/- 11.9*** | 18 | 74.9 +/- 15.2*** | 21 | 73*** | 15 |
|  | High wsf | 99.6 +/- 11.4 | 15 | 119.9 +/- 11.7 | 15 | 86.1 +/- 8.5 | 19 | 105.6 +/- 9.0 | 19 | 16.8 +/- 7.0 | 15 | 18.4 +/- 4.8 | 19 | 25.9 +/- 7.1*** | 20 | 16.3 +/- 6.6 | 20 | 91.7 +/- 5.6 | 20 | 0 | 20 |
| Late exposure 3 dph | Control | 98.0 +/- 7.8 | 50 | 122.6 +/- 10.8 | 50 | 85.7 +/- 16.3 | 61 | 102.9 +/- 19.2 | 61 | 19.7 +/- 6.3 | 50 | 16.2 +/- 6.6 | 61 | 12.2 +/- 8.2 | 79 | 25.8 +/- 6.8 | 79 | 83.2 +/- 7.7 | 59 | 2 | 61 |
|  | Low dose | 99.3 +/- 10.3 | 37 | 122 +/- 13.1 | 37 | 83.1 +/- 10.1 | 49 | 102.2 +/- 12.9 | 49 | 17.8 +/- 8.2 | 47 | 17.0 +/- 5.4 | 47 | 12.0 +/- 6.8 | 59 | 25.1 +/- 5.8 | 59 | 82.9 +/- 4.7 | 77 | 0 | 65 |
|  | Medium dose | 97.6 +/- 12.3 | 40 | 120.9 +/- 14.0 | 40 | 83.0 +/- 16.0 | 50 | 100.2 +/- 19.3 | 50 | 18.6 +/- 7.5 | 40 | 16.7 +/- 7.9 | 50 | 17.2 +/- 9.4** | 58 | 22.1 +/- 8.3** | 58 | 83.4 +/- 5.0 | 60 | 4 | 50 |
|  | High submerged | 97.9 +/- 9.5 | 57 | 121.8 +/- 18.4 | 57 | 75.4 +/- 12.3*** | 69 | 83.1 +/- 15.3*** | 69 | 18.8 +/- 8.0 | 57 | 9.9 +/- 7.0*** | 69 | 23.4 +/- 11.7*** | 78 | 23.5 +/- 6.4 | 78 | 82.8 +/- 4.5 | 78 | 9* | 69 |
|  | High surface | 97.5 +/- 9.9 | 19 | 118.8 +/- 14.6 | 19 | 69.6 +/- 7.6*** | 20 | 71.3 +/- 9.7*** | 20 | 17.5 +/- 7.4 | 19 | 2.3 +/- 3.4*** | 20 | 28.3 +/- 8.2*** | 20 | 21.7 +/- 3.7 | 20 | 84.8 +/- 4.8 | 19 | 60*** | 20 |
|  | High wsf | 99.9 +/- 10.0 | 18 | 124.4 +/- 8.9 | 18 | 84.1 +/- 9.7 | 21 | 102.2 +/- 11.3 | 21 | 19.7 +/- 6.2 | 18 | 17.6 +/- 5.2 | 21 | 23.0 +/- 8.4*** | 20 | 16.3 +/- 2.8*** | 20 | 90.0 +/- 4.6*** | 19 | 0 | 21 |

#### 3.5.2. Functional abnormalities

In both exposures ventricular contractility (VFS) was reduced (Table 2). Notably, reduced contractility was also evident in the lower doses in the *early* exposure. In addition, several animals suffered from a non-contracting or constantly contracting ventricle, defined as silent ventricle (SiV). In the *early* exposure, we observed a dose-dependent prevalence of SiV in all groups with oil droplets, with the most occurrences (73%) in the *H surf* group with the severely abnormally shaped hearts. The only group in *early* exposure with no incidence of SiV was *H WSF*. In the *late* exposure, there were fewer SiVs in all doses, however, we still observed SiV in 60 % of the *H surf* animals, even though their hearts’ morphology looked normal compared to the *early* exposed animals. Reduced heart rate (bradycardia) was only observed in the high surface at the *early* exposure, while increased heart rate (tachycardia) was seen in *H WSF* in the *late* exposure (Table 2).

Edema formation, a consequence of circulation abnormalities, was seen among more treatments in the *late* exposure (Table 2). In the *early* exposure, both *H WSF* (26%) and *H surf* (30%) group had increased frequency of edema compared to the control group (15%). While in the *late* exposure, medium (17%) and all high doses (*H WSF* (23%), *H sub* (23%) and *H surf* (28%)) all showed increased frequency of edema compared to control (12%) (Table 2). Yolk sac size is dependent on hatching time and rate of yolk consumption. We observed an increased average yolk sac size, indicating a decreased yolk utilization in the *H surf* group after *early* exposure. The opposite was observed in the *late* exposure, a decreased yolk sac, thus an increased yolk utilization rate in the *M* and *H WSF* groups (Table 2).

### 3.6. Temporal expression of genes in response to *early* and *late* oil exposure

Generally, the response in gene expression was different for each gene between the two exposures (Fig. 7). Oil-induced differential expression of *bmp10* was independent of circulation with an immediate trend of up-regulation in the *late* exposure. While *rhag* and *abcb1* were delayed, with expression increasing after *cyp1a* expression had increased. Expression dynamics for the control group in all genes are shown in Fig. S8.

**Figure 7:**
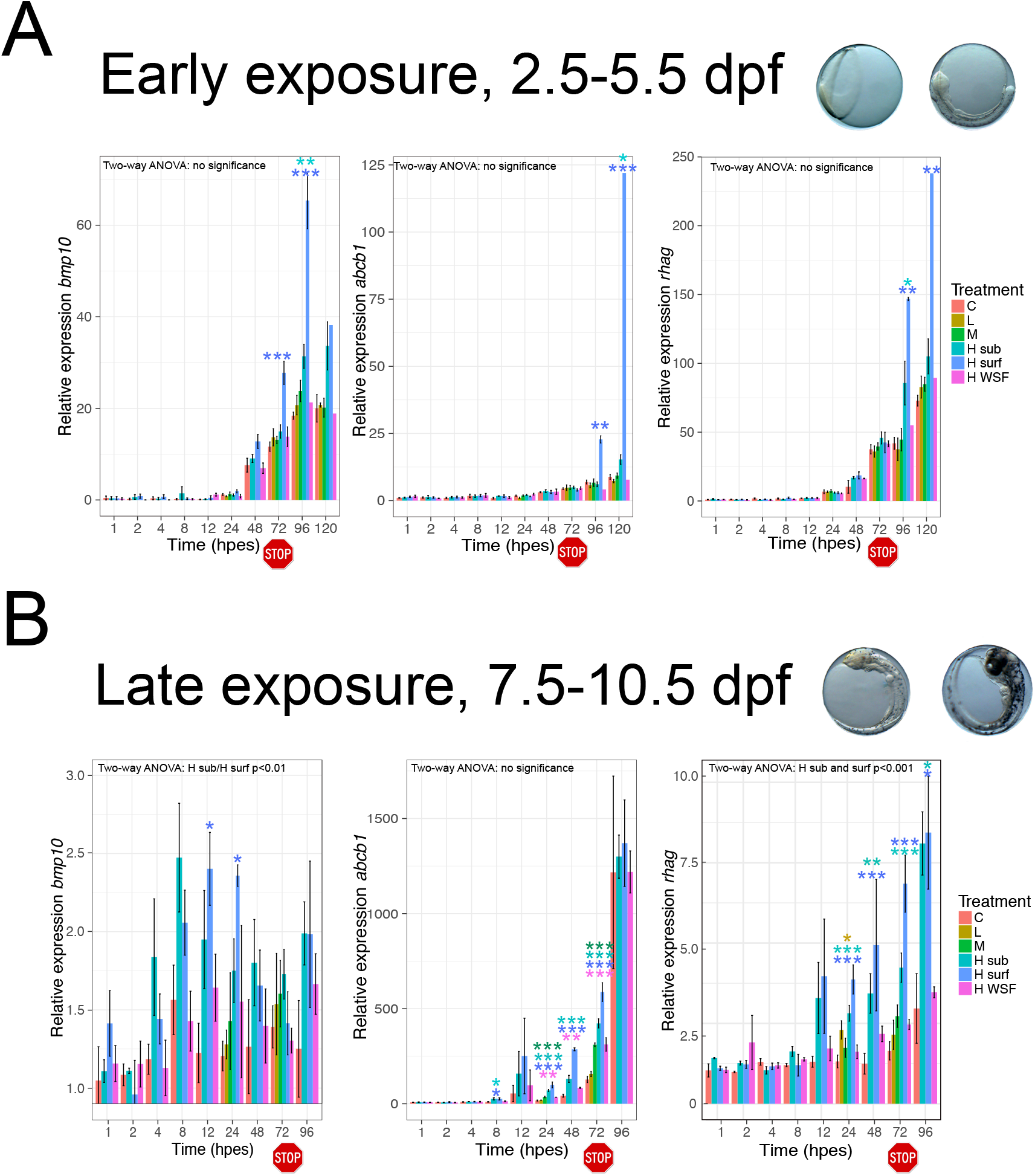
Relative expression of bmp10, abcb1 and rhag. Relative expression compared to control at exposure start for bmp10, abcb1 and rhag after early (A) and late (B) exposure. Expression was related to control group at exposure start to detect both developmental and treatment related expression. Exposure stop is indicated by a stop icon. Statistical difference to control was tested at each time point with one-way ANOVA with Dunnet’s multiple comparsions, and significant differences are indicated by *=p<0.05 or ***: p<0.001. At 120 hpes in the early exposure, only one sample was collected for H surf and H WSF.Dpf: days post fertilization, hpes: hours post exposure start, C: control, L: low dose, M: medium dose, H sub: high dose submerged, H surf: high dose surface, H WSF: high dose water soluble fraction.

Exposure-related up-regulation of *bmp10*, a calcium regulated gene involved in cardiogenesis, was observed before first heart beat at the cardiac cone stage (72 hpes), 2.4 fold expression in *H surf* compared to control (Fig. 7). The exposure start in the *early* exposure was at the beginning of gastrulation, prior to or at the time of onset of zygotic transcription. Thus, at exposure start no expression of the potent signaling molecule, Bmp10, was detected at the mRNA level. Induction of *bmp10* (~7 fold) was first initiated at 4 dpf (48 hpes) in all groups including control and showed an increasing expression throughout the measured period. Significant up-regulation in the *H surf* group was observed at 72 and 96 hpes in the *early* exposure (Fig. 7A). At the *late* exposure the expression of *bmp10* was more variable throughout the period. A trend of up-regulation was seen from 4 hpes. In particular, *H sub* and *H surf* groups were significantly different from control overall in the investigated period (Two-way ANOVA p<0.01) (Fig. 7B). No significant expression changes were detected at 3 dph after either early or late exposure (Fig S7E).

Significant up-regulation of the mRNA encoding the transporter protein Abcb1 was seen after *early* exposure but during *late* exposure (Fig. 7). In the *early* exposure, up-regulation was only seen in *H sub* and *H surf* groups (Fig. 7A). While in the late exposure, up-regulation was observed in medium and all high doses (Fig. 7B). No differential expression of *abcb1* was seen at 3 dph after any exposure treatment (Fig. S7H).

Up-regulation of the mRNA encoding the ammonium and carbon dioxide transporter Rhag, was seen in *H sub* and *H surf* groups at both exposures (Fig. 7). In the *early* exposure, an oil induced up-regulation of 3.5 fold (H surf) and 2 fold (H sub) was observed first at 96 hpes. Differential expression of *rhag* was much more immediate in the *late* exposure (after completion of organogenesis), and significant up-regulation was seen in both *H sub* (2-3 fold) and *H surf* (3-4 fold) groups at 24 hpes and throughout the examined period. Additionally, low dose up-regulation at 24 hpes (Fig. 7B). No differential expression of *rhag* was detected at 3 dph in either *early* or *late* exposure (Fig. S7G).

#### 3.6.1. Whole mount in situ hybridization of rhag

To determine location of *rhag* in early life stages of fish, we performed whole mount in situ (WISH) on unexposed 10 dpf embryos (Fig. 8). In unexposed 10 dpf Atlantic haddock embryos, expression of *rhag* was observed in a line following the branchial arch blood vessels (Fig. 8). No expression was found anywhere else in the embryos, however, due to auto staining, the yolk sac was removed prior to coloration, and thus, the syncytial layer and yolk sac were not available for analysis. The technical negative control (sense) showed no coloration (Fig. 8C).

**Figure 8.**
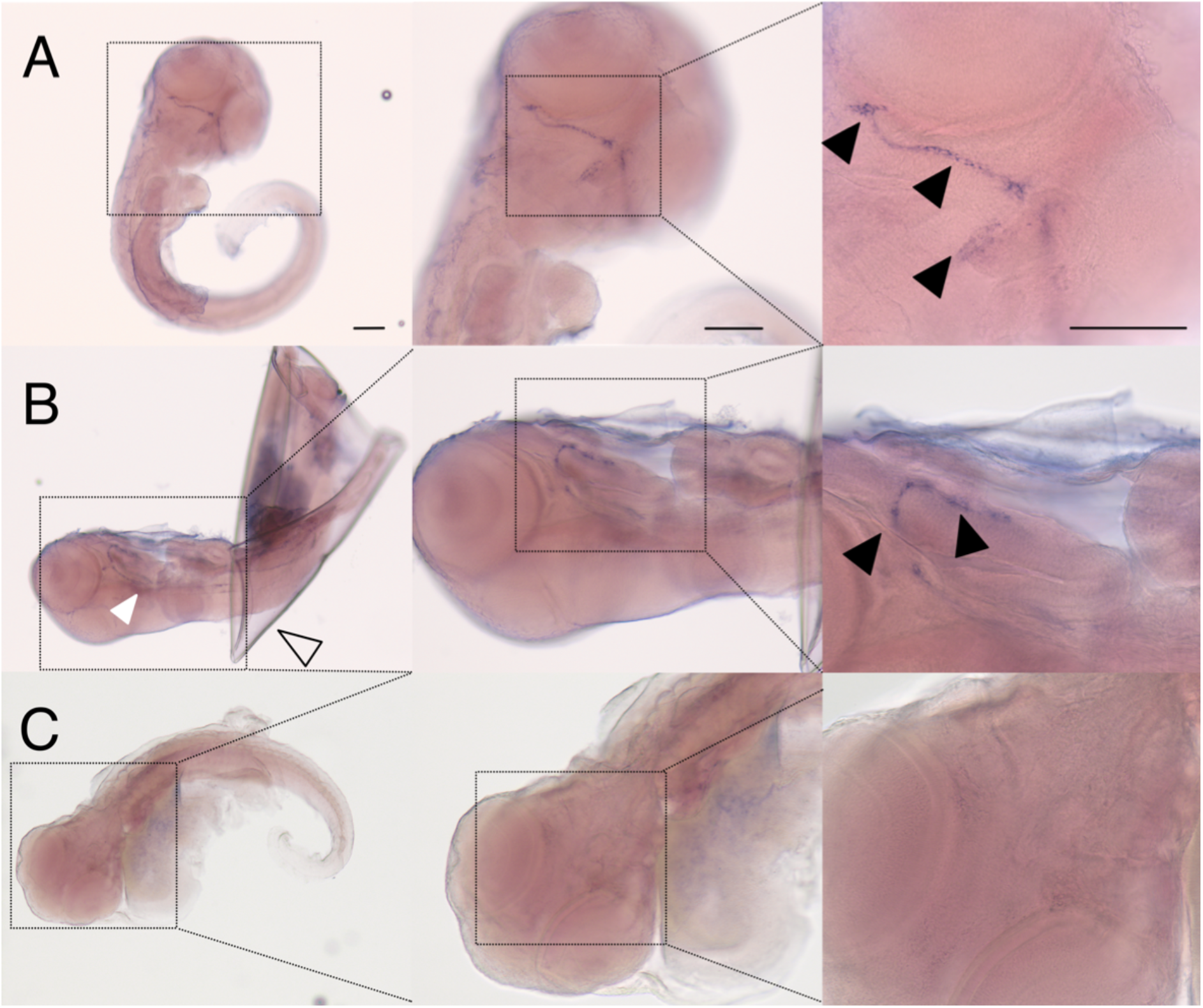
Whole mount in situ hybridization of rhag. Expression pattern of rhag in ventral (A) and lateral (B) view in 10 dpf unexposed embryos of Atlantic haddock. Expression is following the branchial arch. C) No labeling in sense control. Black arrow head: expression of rhag, open arrow: eggshell, white arrow head: unspecific coloration of otholites.

## 4. Discussion

In this study we aimed to link the consequences of short-term crude oil exposures in two embryonic phases to metabolism and abnormal circulation by investigating morphological and functional phenotypes and related gene expression. In general, the embryotoxic consequences of an oil exposure were more severe in the *early* exposure animals. Severe cardiac and craniofacial abnormalities in the highest doses exposed to micro oil droplets were highlighted. The large difference in phenotypic outcome in *early* vs *late* exposure could be explained both by the prolonged exposure and immature response system. First, particularly in the *early* exposure, the groups exposed to micro oil droplets experienced a prolonged exposure evident by visible oil droplets on the chorion and a prolonged *cyp1a* response. Second, the *early* exposure started prior to or at the time of onset of zygotic transcription, in which the embryo has a less efficient response and detoxification system (Sørhus et al., 2016a). Consequently, we observed a delayed response in gene expression of all genes evaluated, including *cyp1a* which is crucial for the first phase in xenobiotic metabolism (Goksøyr, 1995). Higher oil droplet binding and most likely, accumulation, contributed to a higher body burden of PAHs in the *early* exposure, while the levels of PHE metabolites were 2-3 times higher in the *late* exposure. These results are in line with studies showing that Cyp1a induction has a protective rather than aggravating effect (Incardona et al., 2005; Wells et al., 1997; Wells et al., 2009).

Exposing embryos at two developmental time points gave us a unique opportunity to phenotypically anchor temporal gene expression changes and continue to unravel underlying mechanisms of crude oil toxicity. Exposure during early cardiac development, and after onset of heartbeat enabled us to evaluate whether the expression response for the various genes were due to direct effect or secondary to abnormal circulation. Furthermore, we linked gene expression to metabolism by investigating expression in context of Cyp1a regulation, PAH uptake and appearance of metabolites. Mainly, we suggest that *cyp1a, cyp1b, cyp1c* and *bmp10* are directly induced by parent compounds in the oil. To illustrate, these genes are expressed in the *late* exposure before metabolites are detected. Subsequently, *abcb1* and *rhag*, which are linked to transport of xenobiotic products (Choudhuri and Klaassen, 2006) and nitrogenous waste (Caner et al., 2015), respectively, showed a delayed response corresponding to xenobiotic energy consumption utilizing amino acids (Wang et al., 2019) and transport in xenobiotic metabolism (Kimura et al., 2007).

### 4.1. Oil droplet fouling

Oil droplet fouling occured in both *H surf* and *H sub* groups and consequently increased haddock embryotoxicity by escalating the uptake and prolonging the exposure. Droplet fouling on the submerged exposures show that accumulation of oil droplets was due to the biological characteristics of the haddock embryo and not a surface slick artifact. Unlike other cod fish eggs, haddock have an extra membrane of adhesive material covering the primary egg envelope (Hansen et al., 2018; Morrison et al., 1999; Oppen-Berntsen et al., 1990). The chemical composition of this outer membrane is still unknown. However, it clearly has a hydrophopic interaction with oil, and it is responsible for the accumulation of oil droplets on the eggshell. The outer membrane degrades during the embryo phase, a few days before hatching (Sørhus et al., 2015, 2016). Accordingly, the early egg stages accumulated more oil droplets than the late stages in the present study.

Haddock spawn at depths between 30 and 500 meters (Olsen et al., 2010; Solemdal et al., 1997). The eggs have a natural high buoyancy, and all embryonic stages have been observed in the surface at surveillances (Solemdal et al., 1997). Oil droplet fouling can increase the buoyancy of the egg, both by the low density of oil itself, but also by trapping of air bubbles on the chorion (Sørhus et al., 2015). In contrast, we have also observed that haddock eggs with oil droplets become negatively buoyant just before hatching and then sink to the bottom of the exposure tanks. Similar effects have been observed in oil-exposed mahi mahi (*Coryphaena hippurus*) (Pasparakis et al., 2017). The mechanism behind this remains unclear, but one possibility is loss of water due to disruption of the perivetiline membrane or disruption of the osmotic control (Pasparakis et al., 2017). Even though we did observe some slick formation as seen in the *H surf* group, we argue that this group is also the more ecologically relevant exposure dose. In a situation where crude oil is discharged at depths like in the Deepwater Horizon accident or in a rough sea, oil droplets will be located throughout the water column (Hansen et al., 2019; Nordtug et al., 2011). Consequently, oil droplet fouling increases buoyancy, that will lead to increased speed to the surface and to even greater exposures in the oil slick.

Oil droplet fouling most likely causes transfer of more and higher molecular weight oil components to the embryo. Although the nominal oil water dose in *H WSF* were higher than *M* sub group, both PAH uptake and *cyp1a* induction were mainly higher in *M* sub group compared to *H WSF* group especially in the *early* exposure. Interestingly, the composition of PAHs was also different. Nearly no high molecular weight (5-6-rings) PAHs were found in the PAH tissue of the *H WSF* group in any exposure. Accordingly, the *H WSF* group had very little Cyp1a activity (EROD assay) compared to all other treatments where oil droplets were included.

In haddock embryos, oil droplets possibly add a dual toxicity. First, oil droplet fouling results in an increased local concentration of PAHs and other toxic oil components. Second, oil droplets adhered to the chorion may also contribute to transfer of lipophilic compounds that are less available in a general water exposure (Sørensen et al., 2017). The latter may be linked to the high molecular weight PAHs which were not present in the *H WSF*. Enhanced toxicity after direct contact with oil and accumulation on the chorion have also been observed in medaka (*Oryzias latipes*) (Gonzalez-Doncel et al., 2008) and polar cod (Laurel et al., 2019), together with our results it highlight the importance of including dispersed oil in experimental exposure designs and risk assessment models (Nordtug et al., 2011).

### 4.2. PAH uptake, Cyp induction and metabolism

We observed an immature detoxification system in early embryos via delayed expression of *cyp1* paralogs and *abcb1* transporter, higher body burdens of PAHs and lower levels of metabolites. Xenobiotic metabolism is necessary to excrete toxic substances. Among the thousands of potential PAH metabolites, we used PHE as a model compound to study the metabolic capacity of the haddock embryo. Even though PHE is a weak inducer of *cyp1s*, we report PHE metabolite levels as to represent metabolite formation of any kind. The *late* exposure had the more efficient elimination of PAHs evidenced by the immediate strong *cyp1* response resulting in lower peak levels of tissue PAHs and higher levels of metabolites (e.g. PAH levels were approximately 2.5 times lower in *H surf late* compared to *early* exposure). In the *late* exposure *cyp1a, cyp1b* and *cyp1c* mRNA was highly and quickly induced, and already after 24 hours the metabolic capacity surpassed the uptake speed of PHE. This resulted in a reduced body burden even under continuous exposure. However, we did not observe any high accumulation of PHE metabolites, showing that the embryos can effeciently excrete metabolic products during phase III metabolism.

Cyp1a mRNA was clearly the first and the main detoxification enzyme mRNA in *early* embryonic development. The other *cyp1* paralogs, *cyp1b* and *cyp1c*, were not induced until later in the *early* exposure. *cyp1b* was upregulated from 3.5 dpf (24 hpes) regardless of exposure including control and emphasizes the other non-detoxifying roles of Cyp1s (e.g. steroid biosynthesis and cardiovascular and eye development (Chaudhary et al., 2009; Li et al., 2017)) during early fish development (Goldstone et al., 2010). At 72 hpes a 200 fold developmentally related up-regulation of *cyp1c* was observed and at the same time a strong up-regulation was seen in the treatment groups. These observations suggests that xenobiotic induction of *cyp1c* is occurring as soon as the organism possess Cyp1c acitivity. A delayed expression of *cyp1b* and *cyp1c* underscore the differences in capability for fish embryos to metabolize xenobiotics depending on their embryonic developmental window. Our results coincide with the findings of (Kuhnert et al., 2017) where the expression of *cyp1b* and *cyp1c* is also delayed in zebrafish exposed to benzo(a)pyrene.

While Cyp1a appears protective and is necessary for detoxification (Billiard et al., 2006; Mu et al., 2016; Scott et al., 2011; Wincent et al., 2015), it may also result in the production of some reactive toxic metabolites (Wells et al., 1997). Other enzymes can also produce metabolites that can increase the toxicity (Nebert et al., 2004). Several studies have found that hydroxylated PAHs have greater toxic effects than the parent compounds (Chibwe et al., 2015; Diamante et al., 2017; Fallahtafti et al., 2012; Schrlau et al., 2017). Bioactivation of PAHs has also been linked to endocrine disruption (Fernandes and Porte, 2013; Hyzd’alova et al., 2018; Pencikova et al., 2019; Sievers et al., 2013; Van de Wiele et al., 2005). Moreover, electrophilic intermediates of PAH metabolism are well known for the formation of hydrophobic DNA adducts and cancer induction (Moorthy et al., 2015). PAH metabolites may also play an important role in cardiotoxicity. PHE and oil are shown to block currents of K^+^ through the ERG potassium channel (Brette et al., 2014; Brette et al., 2017). However, the exact mechanism behind how three-ring PAHs are able to block the channel is not well defined (Incardona, 2017). *In silico* docking models suggest that PHE has the necessary three aromatic rings, but lacks any other functional groups essential for strong blocking capacity (Cavalli et al., 2012; Vandenberg et al., 2012). Other types of membrane ion channels have been intensively studied through the use of PAH metabolites: The 9-hydroxy-PHE metabolite (9-phenanthrol) is a selective and sensitive blocker of the transient receptor potential melastatin 4 channels (TRPM4) (Burris et al., 2015; Guinamard et al., 2014; Hou et al., 2018) and anthracene-9-carboxylic acid is a commonly used blocker of the calcium-activated chloride channels (CaCCs) (TMEM16A) channels (Cherian et al., 2015; Ta et al., 2016). The blocking capacity of PAH metabolites to the ERG channels has not yet been studied to our knowledge, but it is possible, if not likely, that critical aspects of crude oil cardiotoxicity are caused by an ERG-blocking PAH metabolite. Taken together we see a need for more studies of the effects of PAH metabolism in oil toxicity, and urge more labs to develope analytical methods able to cover a broad spectra of oil-related metabolites. The latter is a great methological challenge due to extreme complexcity in the number of metabolites and the lack of available standards.

We show that *cyp1* mRNA induction, PAH tissue content, and PAH metabolites are related to early life stage toxicity in fish, where the amount of mRNA induction and PAH and metabolite levels is related to the window of exposure. This observation is in line with an established relationship between toxicity and PAHs in crude oil (Carls and Meador, 2009; Hodson, 2017) and petrogenic substances (Adams et al., 2014; Kamelia et al., 2019; Kamelia et al., 2017). However, there are multiple toxic components in crude oil apart from PAHs that can induce toxicity in early life stages of fish (Meador and Nahrgang, 2019; Sorensen et al., 2019). Therefore, there is a need for more fractionation studies and Effect Directed Analysis (EDA) to increase our understanding of the additive and synergetic effects of the complex oil toxicity.

### 4.3. Morphological abnormalities

The large differences in morphological abnormalities after *early* vs *late* exposure, show the importance of timing of exposure. The *early* exposure has major consequences for formation of organs by interfering with the developmental signaling pathways, while *late* exposure mainly shows effect on organism condition possibly downstream of cardiofunctional defects.

The eye abnormalities were most likely downstream of both circulation defects and altered cyp-balance. Crude oil-induced eye developmental defects are well documented. Both reduced eye size (anapthalmia) and reduced size with abnormalities (microphtalmia) are observed (Incardona, 2014; Lie et al., 2019; Magnuson et al., 2018). In the present study, eye size was mainly affected in the *early* exposure, where we find the most severe developmental and functional abnormalities. Zebrafish with no cardiac function (*tnnt* morpholinos) mimic the oil-induced reduction in eye size (Incardona et al., 2004), suggesting that anapthalmia is secondary to circulatory defects. However, it is also suggested that abnormal cyp-balance disrupts retinoic acid signaling pathway through disturbance of Cyp26, leading to abnormal eye development (Lie et al., 2019; Yamamoto et al., 2000). Another Cyp enzyme, Cyp1b, is also involved in eye development (regulating ocular fissure closure) through both retinoic acid-dependent and -independent pathways (Chambers et al., 2007; Williams et al., 2017). Affirmative of a non-toxicological function of Cyp1b was the general increase in expression of *cyp1b* beginning at 3.5 dpf (24 hpes). Overexpression of *cyp1b* causes irregular eye shape referred to as coloboma (Chambers et al., 2007; Williams et al., 2017). The *late* exposure animals possessed less severe abnormalities in general. However, the *H sub* and *H surf* groups still showed excessive incidence of eye abnormalities (irregular eye shape) which were reflected in increased *cyp1b* expression. These findings are in line with findings that proper closure of the ocular fissure is disrupted by overexpression of *cyp1b* (Chambers et al., 2007; Williams et al., 2017).

The craniofacial abnormalities observed in our exposed animals are potentially due to both disruption of jaw and pharyngeal arch formation and reduced outgrowth linked to abnormal craniofacial muscle function. Expression of Cyp1b was found in pharyngeal arches of zebrafish exposed to PCB 126 and TCDD (Timme-Laragy et al., 2008; Yin et al., 2008), and the injection of human *cyp1b* mRNA resulted in disruption of neural crest-derived jaw and pharyngeal arch formation (Williams et al., 2017). Furthermore, muscle contraction is mandatory for proper stacking of chondrocytes and outgrowth of craniofacial cartilages (Shwartz et al., 2012). Crude oil disrupts cardiac muscle function (Brette et al., 2014; Brette et al., 2017; Incardona, 2014), and jaw spasms observed in previous studies suggest a similar effect on the craniofacial muscles. Impairment of craniofacial muscles might explain the high incidences of craniofacial abnormalites in the *late* exposure. In conclusion, the tight correlation between *cyp1b* expression and oil induced eye and jaw deformities suggests that inappropriate expression of *cyp1b* in oil-exposed animals are directly linked to abnormal eye and jaw development.

### 4.4. Cardiac morphological defects

The severely malformed and small hearts after *early* exposure imply multiple impacts on cardiac formation and early development, while the reduced heart size after *late* exposure suggests on late development and especially cardiomyocyte proliferation.

A circulation- and time-independent expression of *bmp10* suggests a direct effect of crude oil on calcium homeostasis. Several signaling pathways linked to cardiac development and proliferation are impacted by the complexity of crude oil exposure, including calcium-regulated pathways (Sørhus et al., 2017; Xu et al., 2017), AhR (Yin et al., 2008) and retinoic acid (Lie et al., 2019)-dependent pathways. Disruption of intracellular calcium levels has been shown to reduce cardiomyocyte proliferation leading to smaller ventricles (Ebert et al., 2005; Rottbauer et al., 2001), suggesting that ventricular proliferation is calcium controlled. Bmp10 is a calcium-regulated signaling molecule involved in cardiogenesis (Huang et al., 2012; Wamhoff et al., 2004; Wamhoff et al., 2006) and is thought to orchestrate several major key cardiogenic factors and play a part in ventricular cell proliferation (Chen et al., 2004; Shou et al., 1998). In the current study, we observed an up-regulation of *bmp10* regardless of timing of exposure. However, the developmental consequences of timing of an inapropriate gene regulation is essential. In the *early* exposure, the initiation of *bmp10* up-regulation was occurring at cardiac cone stage (72 hpes). At this developmental time point asymmetric Bmp signaling (in particular bmp4) in the cardiac primordium is involved in the looping of the heart (Chocron et al., 2007; Huang et al., 2012; Lombardo et al., 2019). An overexpression of *bmp10* in the cardiac primordium may therefore disrupt left/right looping of the heart. At the start of the *late* exposure, the looping process was already initiated, and the heart was functionally beating. Consequently, we observed less severe effect on looping, while growth of the heart was altered. Reduced size of the ventricle was observed in both *H sub* and *H surf*, as well as an overall up-regulation of *bmp10*. The up-regulation of *bmp10* may be consequential for the reduced ventricle size in exposed larvae from the late embryonic stages. Taken together, the observations from *early* and *late* exposures support that the oil components have a circulation-independent direct effect on calcium homeostasis likely disrupting cardiac development and proliferation indicated by the up-regulation of *bmp10*.

### 4.5. Cardiac functional defects

The cardiac functional defects post-exposure were linked to irreversible morphological changes and reversible effects of circulating crude oil components. Reduced contractility and non-contracting or constantly contracting ventricles (SiV phenotype) were traits found in both *early* and *late* exposures. Functional defects in the *early* exposure could originate from irreversible morphological changes (severly malformed ventricles) but could also stem from consumption of oil components stored in the yolk sac after hatching as observed in previous studies (Sørhus et al., 2016b; Vanleeuwen et al., 1985). The *late* exposure animals, on the other hand, had normally shaped but often smaller ventricles. The contractility abnormalities most likely stem from components stored in the yolk sac.

A suite of defects downstream of cardiac dysfunction has been described for oil-exposed developing fish, including edema accumulation (Incardona, 2017; Incardona and Scholz, 2016). Edema formation measured included both pericardial and yolk sac edemas, and may be due to i) irreversible morphological cardiac deformities that alter the cardiac function later in life and ii) oil components that can directly alter the electrophysiological activity (contractility and rhythm) of the developing heart (Brette et al., 2014; Brette et al., 2017; Incardona, 2017). Circulatory defects may also affect distribution of lipids from the yolk which in turn may affect growth in later stages. Polar cod exposed to transient low levels of oil induced dysregulation of lipid metabolism and reduced growth in morphologically normal juveniles (Laurel et al., 2019). We observed a reduced yolk utilization in *H surf* in the *early* exposure, suggesting that these embryos were deprived of essential yolk-based lipids. Furthermore, the reduced yolk utilization seems to be associated with heart rate, and thus, dependent on circulation. Accordingly, the treatment groups with tachycardia had highly increased yolk utilization (*H WSF*), while the treatement groups with bradycardia (*H surf*) had severely reduced yolk utilization. Overall, cardiocirculatory defects during embryonic development could be sufficient to impact growth and could be consequential for individual fitness at juvenile and adult stages.

### 4.6. Osmoregulation

Up-regulation of *rhag* suggests that osmoregulatory cells are impacted by oil exposure. Osmoregulation can be disrupted by circulatory defects (Miyanishi et al., 2013), or directly affected by oil components (Sørhus et al., 2017). In adult fish, the main localization for osmoregulation, gas exchange and transport of ammonium are in the gills (Evans et al., 2005) in osmoregulatory cells called mitochondrial rich cells (MRCs) (Evans et al., 2005; Hiroi et al., 2005; Hirose et al., 2003; Shelbourne, 1957). In 10 dpf haddock, expression of *rhag* was found in the branchial arches (primordial gills), in close proximity to expected localization of MRCs (Zimmer et al., 2014; Zimmer et al., 2017). This finding coincides with expression of *rhag* in mahi mahi at approximately the same developmental time point (Wang et al., 2019). In both *early* and *late* exposures, *rhag* seemed to track more with the peak of *cyp1a* induction and *abcb1* transporter. This could indicate that energy for Cyp1a-mediated PAH metabolism is derived from amino acids (Pasparakis et al., 2016), generating nitrogenous waste also suggested by Wang et al. (2019). Thus, this increased amount of ammonium waste can also affect osmoregulation capacity of MRCs and consequently lead to increased edema formation. Additionally, ammonia excess in itself can add to the toxicity. In mouse, ammonium perturbs glial cells ability to remove potassium and are therefore toxic to the brain (Thrane et al., 2013). Consequently, the crude oil exposed embryos could experience an internal synergy between internal ammonium toxicity and PAH toxicity.

## 5. Conclusion

In this study we showed that oil droplet fouling occurred throughout the whole water column and contributed to increased PAH uptake and embryotoxicity. Oil droplet fouling makes the haddock embryo extremely vulnerable to even short, low level oil exposures. The high bioaccumulation of PAHs and lower levels of metabolites in the *early* exposure reflects an immature response system that possibly has an exacerbating toxic effect in early embryos. Short term exposures prior to and after onset of heart beat and frequent sampling regime enabled us to establish direct and downstream effects of crude oil exposure. The relatively rapid and circulatory-independent response in *bmp10* regulation suggested direct effects on calcium homeostasis affecting calcium-regulated developmental pathways, including cardiogenesis. Similarly, we propose that abnormal regulation of *cyp1b* due to oil exposure may affect eye and jaw development. Expression of *rhag* indicates a direct oil effect on osmoregulatory cells and osmoregulation. We recognize that oil is a complex mixture of many potential toxic components. Yet, the strong correlations among PAH uptake, *cyp1a* induction, metabolite formation and malformations suggest that PAH tissue body burden remains a good metric for oil toxicity.

Our findings are adding more knowledge about development stage-dependent effects of crude oil exposure. Thus, we provide more knowledge and detail to several existing adverse outcome pathways of crude oil toxicity.

## 6. Acknowledgements

We would like to acknowledge Stig Ove Utskot and Tobias Hukset for breeding and management of the fish, Charlotte Nakken for technical assistance with the metabolite data, Prescilla Perrichon for discussion of the data and review of the manuscript and Karen Peck for a thorough review of the manuscript. This work was financed by the Research Council of Norway (EGGTOX: Unraveling the mechanistic effects of crude oil toxicity during early life stages of cold-water marine teleosts (Project # 267820), www.forskningsradet.no) and the Institute of Marine Research, Norway. The funders had no role in study design, data collection and analysis, decision to publish, or preparation of the manuscript.

## 7. Author contributions

Conceived and designed the experiments: E.S., S.M., Ø.K. and A.T.

Performed the experiments: E.S., S.M., Ø.K.

Analysed and interpreted the data: E.S., S.M., C.E.D., D.D.S.

Contributed reagents/materials/analysis tools: A.T., D.D.S.

Contributed to the writing of the manuscript: E.S., S.M., C.E.D.

## 8. Competing interests

The authors declare no competing financial interests.

## Notes

### Competing Interest Statement

The authors have declared no competing interest.

