## Supplementary Information for "Untangling mechanisms of crude oil toxicity: linking gene expression, morphology and PAHs at two developmental stages in a cold-water fish"

**Localization of gene expression**

Generation of probes for *in situ* hybridization (ISH) to localize expression of Atlantic haddock *rhag* was performed by amplifying cDNA using PCR containing T7 and Sp6 primers in the forward and reverse primers, respectively (Rhag_FW_Sp6: 5′- *TTTAGGTGACACTATAG*CATCCTGTTCGCCTTGTACG-3′ Rhag_RV_T7: 5′- *TAATACGACTCACTATAGGG*GAGTTGAAGCTCGGCCAAAA-3′). The PCR product of 634 bp (CATCCTGTTCGCCTTGTACGCGGAGTATGACGACGGCAAGGGCGACGGCCACGACCCTCACCCCAGCAACTCCACCCCCCACGCCGAGCCGAAGGACCCCATGAGCCTCTACCCC**AT**GTTCCAGGATGTCCACGTGATGATCTTCATCGGCTTTGGCTTCCTCATGACTTTCCTGAAGAGGTACGGCTTCAGCAGCGTGGGCGGTAACCTGCTGCTGGCCGCCTTTGGCCTACAGTGGGGCCTGCTGACCCAGGGCTTCTGGCACCTCACGGACGGCAAGATCAAGATCAACATCTTCAA**GA**TAATCAACGCAGACTTCAGCACAGCCACAGTGCTGATCTCGTTTGGAGCCGTCCTGGGAAAGACCAGCCTGGTCCAGCTGCTCATCATGACCGTGCTGGAGATCGGGATCTTCGCCATCAATGAACACCTGGTGGCTAAAGTATTT**GA**GGCGAACGACGTAGGGGCCTCCATGATCATCCACGCCTTCGGAGCCTACTTTGGCCTGGCGGTCGCTCGGGTTCTGTACCGGCCTGGCCTCAAGAACGGACACGAGAACGACGGCTCCGTCTACCACTCTGATCTGTTTGCCATGATCG**GA**ACCGTCTTCCTCTGGATGTTTTGGCCGAGCTTCAACTC (exon-exon junctions in bold)) was used as a template for synthesizing the sense and antisense cRNA probes by the Sp6 and T7 RNA polymerase, respectively. DIG-AP RNA labeling kit (Roche Molecular Biochemicals) was used to perform digoxigenin (DIG) labeling following the manufacturers protocol.

Eggs were dechorionated in methanol before 5 min rehydration in methanol/1xPBS, 100%, 75%, 50%, 25%. Pigment was removed using 3% H2O2/0.5% KOH medium until complete disappearance of pigmentation, followed by wash in 1xPBS. The embryos were transferred to 125 µm Nylon membranes (Thisse 2007) and proteinase K treated (20µg proteinase K/ml 0,05 TrisHCl pH 7.5) for 10 minutes. After proteinase K treatment, the embryos were post fixated for 20 minutes in 4% PFA, followed by 4x5 minutes wash in PBST (1XPBS 0.1%Tween). The samples were prehybridized in hybridization buffer (50% formamide (Ambion), 5xSSC (Sigma-Aldrich), 5xDenhardts (Sigma-Aldrich), 250 µg/ml tRNA (Roche), 500 µg/ml salmon sperm DNA (Roche), 10% dextran sulfate (Sigma-Aldrich)) for 2 hours at 70°C before hybridization with preheated 600 ng probe/ml hybridization buffer at 70°C for 72 hours. Hybridization was followed by stringent wash steps: 2x15 minutes in 50% formamide in 2xSSC-T (2xSSC (Sigma) and 0.1% Tween 20), 1x30 and 1x15 minutes in 2xSSC-T and finally 2x15 minutes in 0.2xSSC-T. Samples were subsequently treated with 15µg RNAse A/ml RNAse buffer (0,01M tris-HCl, pH 7.5, 0.5M NaCl, 0.005 M EDTA) for 20 minutes at 37°C, followed by wash in RNAse buffer for 20 minutes at 65°C. Antibody incubation was initiated by washing 2x5 minutes in B1 (0.1 Tris-HCl, 0.15M NaCl, pH 7.5, 0.1% Tween 20), before 1-3 hours of incubation in B1 with 2 mg/ml BSA and 2% Heat inactivated goat serum (HING) (B2). Finally, the samples were incubated over night at 4°C in 1:2000 anti-DIG in B2. Before staining the samples were washes 4x1h in B1 and 1mM Levamisole, followed by 2x5 minutes in B3 (0.1M Tris-Hcl, 0.1M NaCl, 0.05 M MgCl2, pH 9.5). Staining was performed by incubation in B3 with NBT/BCIP (338 ng NBT/ml, 169 ng BCIP/ml) and Levamisole (0.24 mg/ml) for 24 hours at room temperature (in dark). Coloration was terminated by adding 4%PFA in PBS before transfer to TEN (10 mM TrisHCl, 1mM EDTA, 0,9% NaCl pH 8.0). Finally, the samples were mounted in glycerol by incubation with 100% glycerol overnight. Microscopical images were taken using a Leica microscope coupled to a Spot Insight cmos camera (Spot Imaging ^TM^).

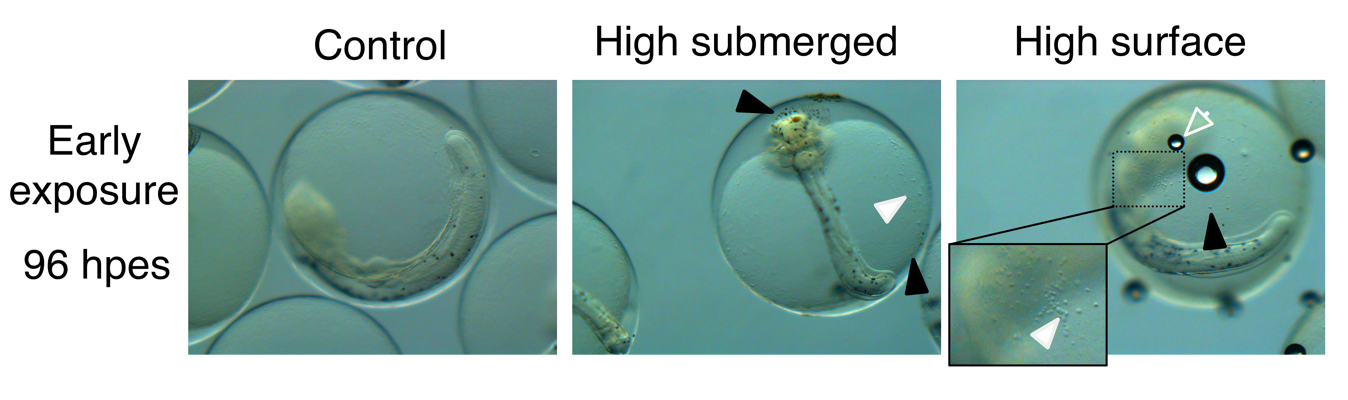

*Supplementary figure S1:* ***Oil droplet fouling 96 hours after exposure start (hpes).*** *Oil droplets on the eggshell 24 hours after exposure stop (96 hpes), some of the oil droplets appear transparent after transfer to clean sea water. Brown (black arrow head) and what appear to be transparent oil droplets (white arrow head) appear on the chorion of both submerged and surface embryos. Open arrow head: trapped air bubble on the chorion. C: control, H sub: high dose submerged, H surf: high dose surface, H WSF: high dose water soluble fraction*

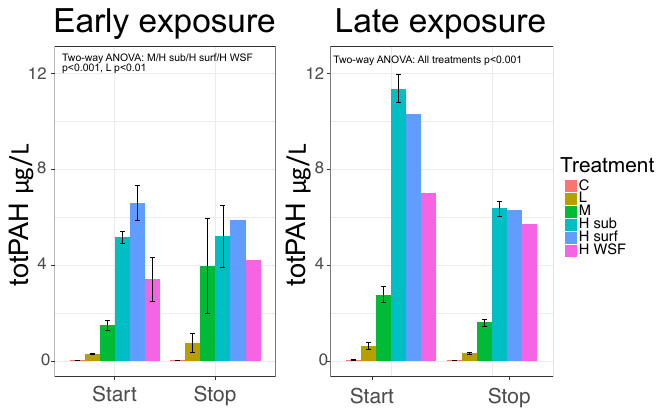

*Supplementary figure S2:* ***Water concentrations of polycyclic aromatic hydrocarbons (PAHs).*** *Total amount of PAH ug/L at exposure start and exposure stop in early exposure, and exposure start, during exposure (24 hours post exposure start (hpes)) and exposure stop in Late exposure. C: control, L: low dose, M: medium dose, H sub: high dose submerged, H surf: high dose surface, H WSF: high dose water soluble fraction. Statistical difference to control was tested with two-way ANOVA with Dunnet´s multiple comparsions, and significant differences are indicated by **=p<0.01 or ***: p<0.001. Note: Only one sample from the H surf and H WSF group was analyzed for end of exposure early exposure and start and end late exposure.*

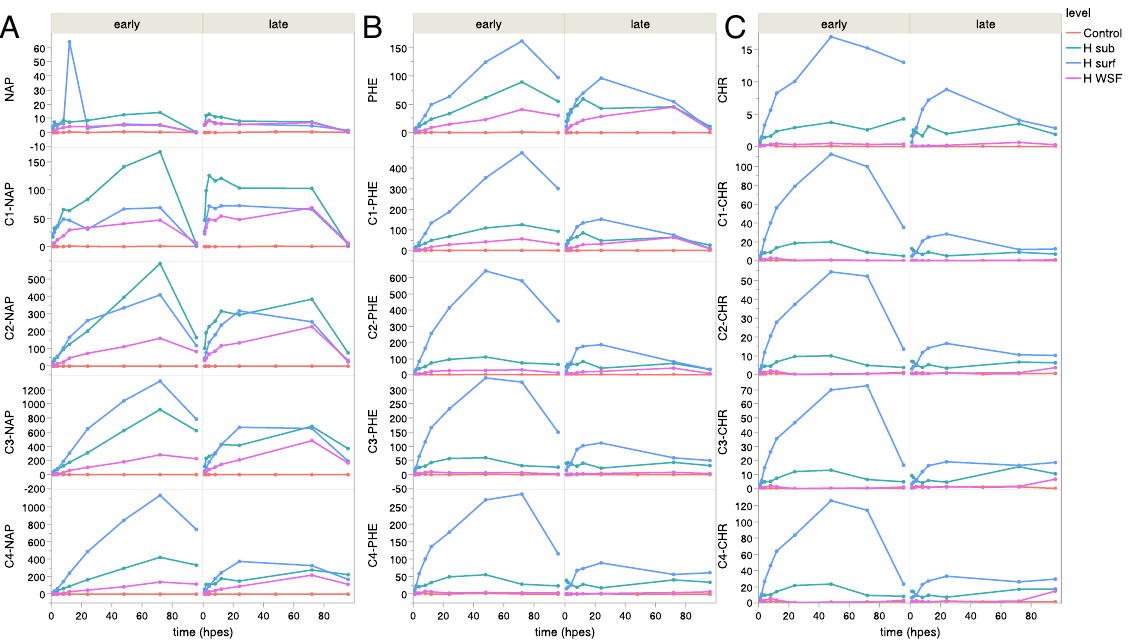

*Supplementary figure S3:* ***PAH tissue burden (ng PAH/g ww) over time (hpes) for classes of PAHs and their alkylated homologues****; A) NAP: naphthalenes, B) PHE: phenanthrenes, and C) CHR: chrysenes. Difference in tissue burden between H surf and the other high dose groups are greatest in higher molecular-weight PAHs or homologues. Hpes: hours post exposure start, C: control, H sub: high dose submerged, H surf: high dose surface, H WSF: high dose water soluble fraction.*

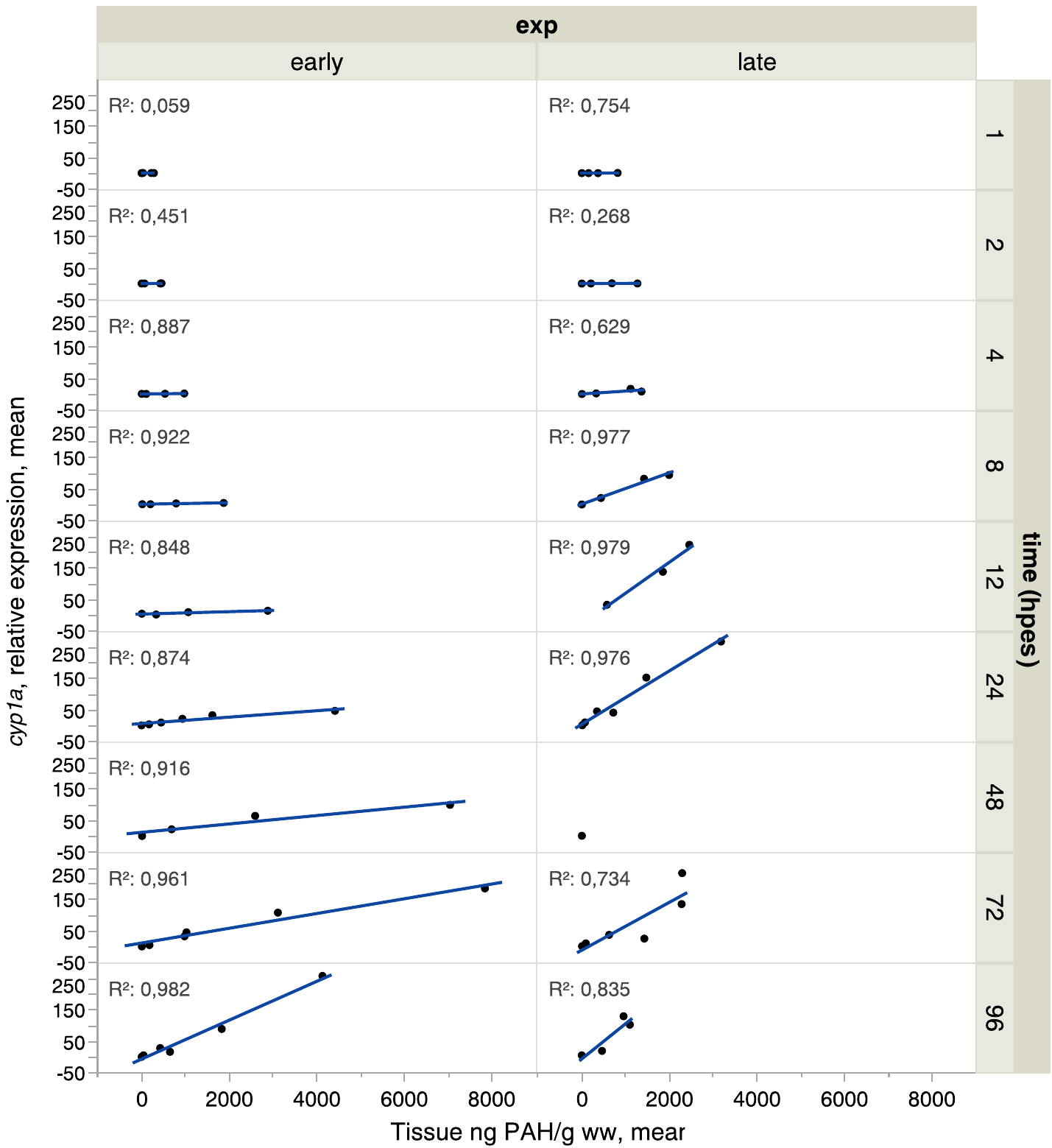

*Supplementary figure S4:* ***Cyp1a expression and PAH tissue burden concentrations*** *are correlated. Relationships are divided by time points (hpes) and early and late experiments. Each point represents the average of all replicates per treatment for both relative expression of cyp1a and PAH tissue burden. R^2^ values are the coefficients of determination of a linear regression. Hpes: hours post exposure start.*

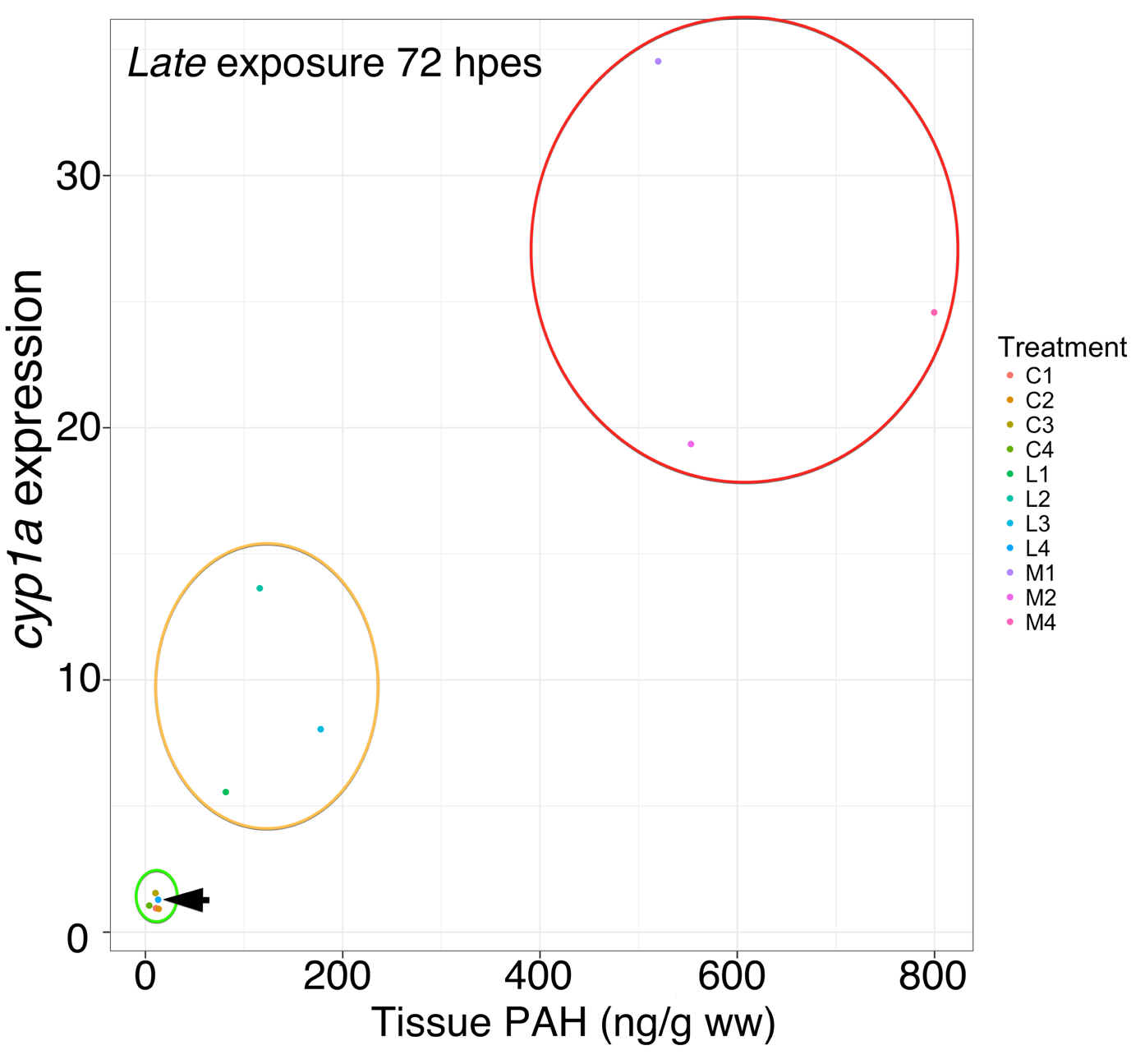

*Supplementary figure S5:* ***Reduced toxicity in outlier tank****. Dose relationship between tissue content of total PAH and cyp1a mRNA levels at 72 hpes in the late exposure. X-axis shows total PAH data with corresponding relative expreesion of cyp1a mRNA levels on the y-axis. The Red circle surrounds medium dose tanks, yellow circle, the low dose tanks (L1, L2 and L3) and green circle the control tanks and low dose tank L4, the latter is indicated by the black arrow. Hpes: hours post exposure start.*

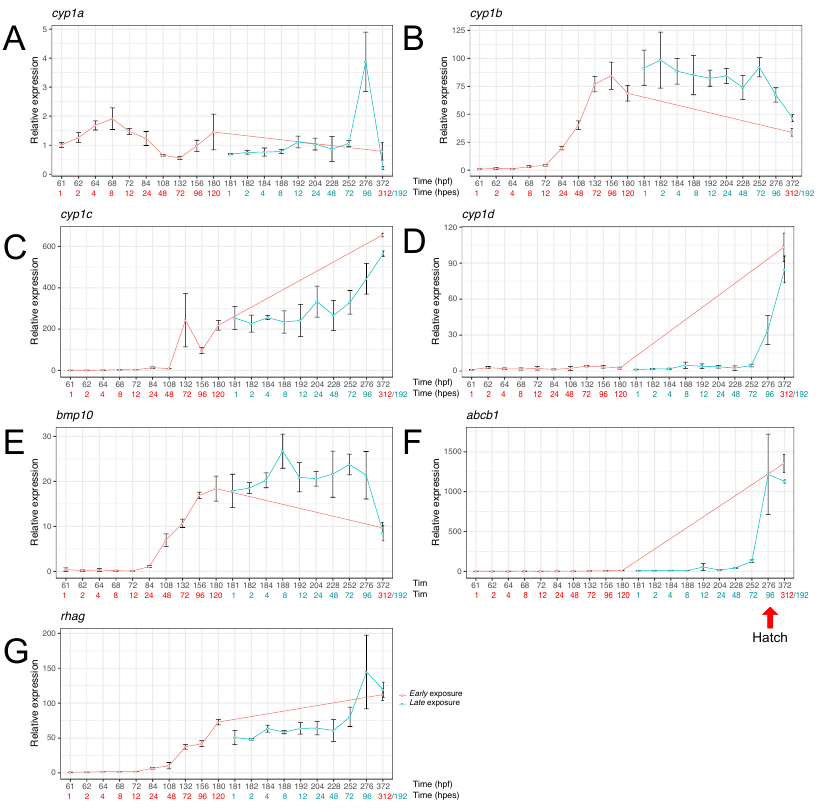

*Supplementary figure S6:* ***Gene expression pattern in control groups in Early and Late exposure.*** *Temporal gene expression in control groups for (A) cyp1a, (B) cyp1b, (C) cyp1c, (D) cyp1d, (E) bmp10, (F) rhag and (****G****) abcb1. 50% hatch was observed at 276 hpf and are indicated with an arrow in (G). Hpf; hours post fertilization, hpes: hours post exposure start.*

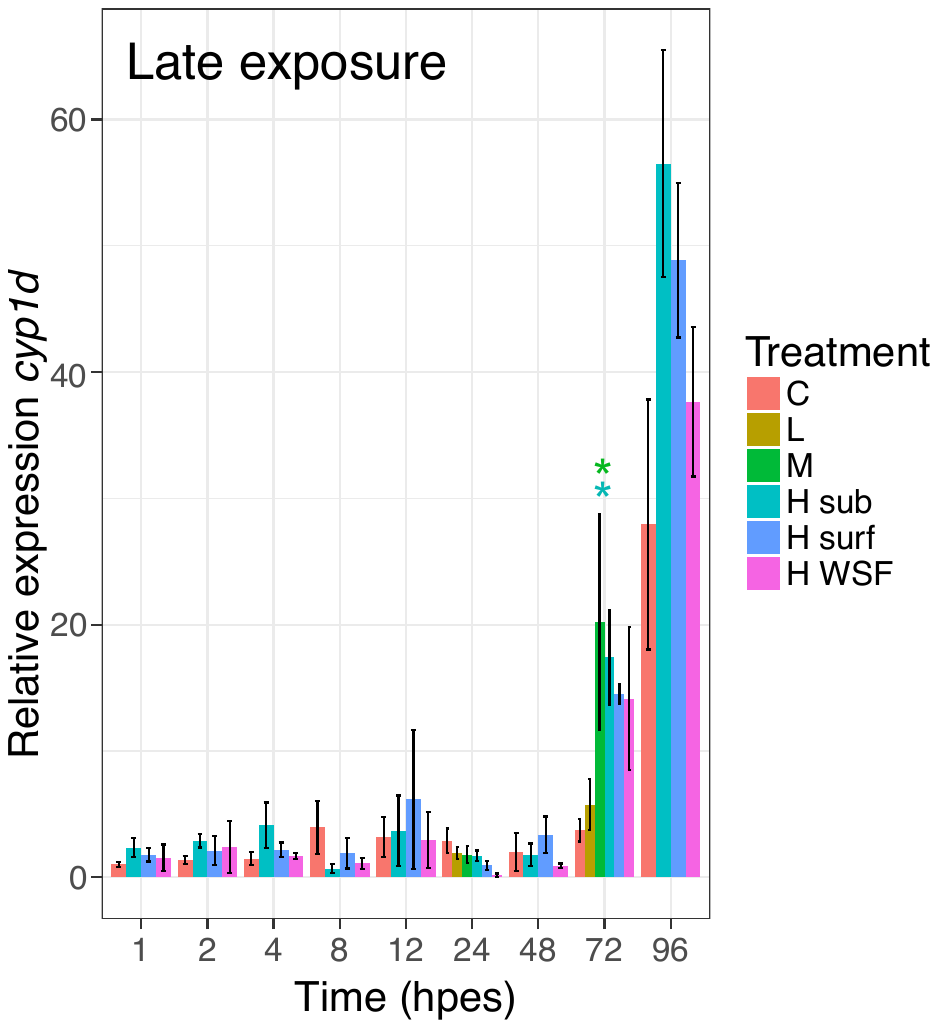

*Supplementary figure S7:* ***Relative expression of cyp1d in late exposure.*** *Expression was related to control group at exposure start. Initiation of cyp1d is observed at hatching stage (96 hours post exposure start (hpes)) in all treatments including control. C: control, L: low dose, M: medium dose, H sub: high dose submerged, H surf: high dose surface, H WSF: high dose WSF. Statistical significant difference from control is indicated by asterisks; p<0.05 *.*

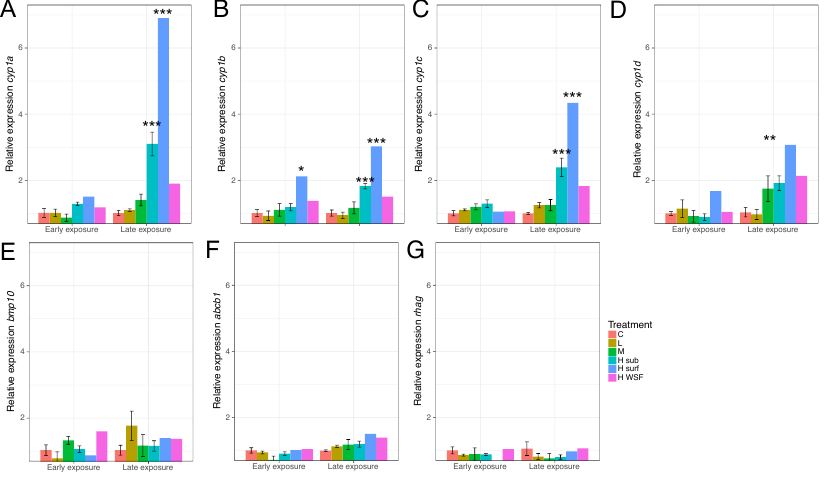

*Supplementary figure S8:* ***Gene expression at 3 days post hatch (dph).*** *Relative expression of cyp1a (A), cyp1b (B), cyp1c (C), cyp1d (D), bmp10 (E), abcb1 (F) and rhag (G) in Atlantic haddock larvae after early and late embryonic exposure. C: control, L: low dose, M: medium dose, H sub: high dose submerged, H surf: high dose surface, H WSF: high dose water soluble fraction. Statistical significant difference from control is indicated by asterisks; p<0.05 *, p<0.001 **. P<0.0001 ***.*

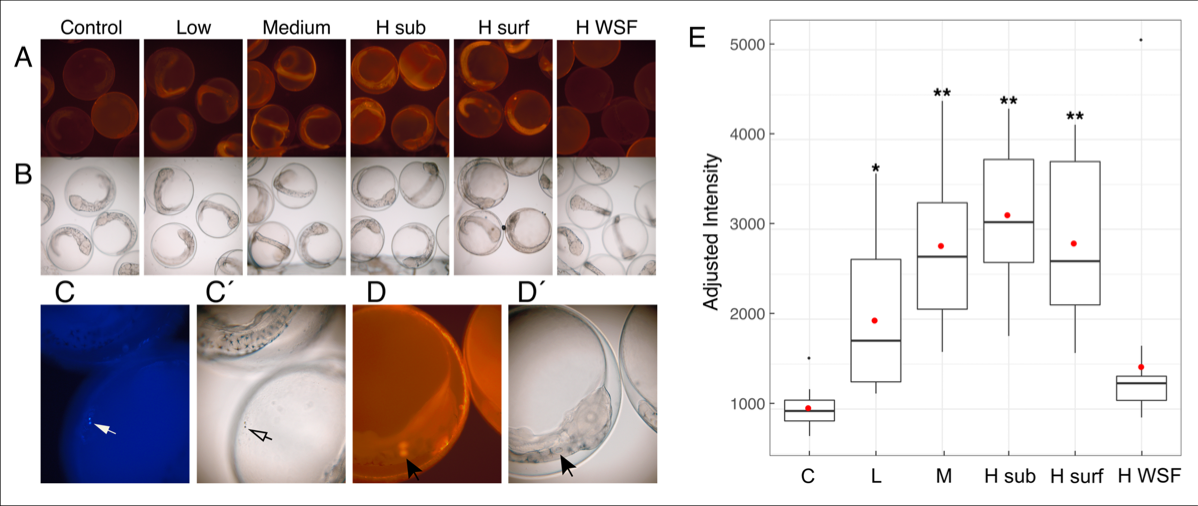

*Supplementary figure S9:* ***Catalytic measurement of Cyp1a by Ethoxyresorufin-O-deethylase (EROD) activity in early exposure.*** *A) Fluorescence (590 nm) images showing EROD staining in control, low dose, medium dose, H sub, H surf and H WSF 72 hours post exposure start (hpes) in early exposure. Staining is mainly apparent in the trunk and otholiths. B) Corresponding brightfield images. C) DAPI (445 nm) image of oil droplets on H sub embryo showing internal fluorescence activity of oil components (white arrow), while C´) shows the corresponding brightfield image, open arrow indicate same oil droplets. D) EROD activity in otholiths (black arrow) in H sub embryo and D´) corresponding brightfield image. E) Fluorescense (590 nm) intensity in all treatments. High dose submerged, H sub; High dose surface, H surf; High dose water soluble fraction, H WSF. Statistical significant difference from control is indicated by asterisks; p<0.05 *, p<0.001 ** (one-way ANOVA with Dunnet´s multiple comparsions test).*

| **Gene** | **Forward** | **Reverse** | **Probe** | **Probe dye** | **Quencher** | **Task** |
| --- | --- | --- | --- | --- | --- | --- |
| *ef1a* | ATCGGCGGTATCGGAACAG | GCTTGAGGACACCGGTCTCA | ACCCGTGGGCCGTG | FAM | none (MGB) | Reference |
| *rxrba* | AAGCAGAAATACCCCGACC | CATGAGGAAGGTGTCGATGG | TGCGTTCCATTGGTCTGAAGTGC | FAM | TAMRA | Reference |
| *abcb1* | CGGGCAGGACAAAGAGATAA | GGTAGATGATCAGGAAGGTGAAG | TGGGCATTAAGAAGGCTGTGTCCA | FAM | TAMRA | Target |
| *cyp1a* | CCTCCTTCCTGCCCTTCAC | TTGGGAATGAAGTAGCCATTGA | CCTCACTGCGCCACAAAAGACACATC | FAM | TAMRA | Target |
| *cyp1b* | GTCTCGGTCAAACAGGACTAC | CCGGAAACCTCACGAGAATTA | CCACTGCGCTTTCGTGGATCATTT | FAM | TAMRA | Target |
| *cyp1c* | GAGCTTGACGTTCACCAATTAC | CTGTCACATGCTGCTCAAAC | TTCCTCTGCCAACAGTGAGACCAA | FAM | TAMRA | Target |
| *cyp1d* | CGTGACCTCCTTCGATAAGAAC | GAATGATCTGGGTGTTGGAGAG | AGGAGAGAGGAGAGGATCAGCATGG | FAM | TAMRA | Target |
| *rhag* | GTCTCTGCTTATGCCATCTCC | CCGAAGGGTCCGATATTCATG | CAGGGTCGCGTTCTGAATGTGGA | FAM | TAMRA | Target |

*Table S1****: Real time qpcr primers and probes.***

*Dataset S1:* ***The PAH tissue (ug PAH/g ww) content at all stages in in early and late exposure.*** *Ww; wet weight, hpes; hours post exposure start, rep; replicate, C0; non alkylated, C1-C4; alkylated homologues BT: Benzothiophene, NAP: Naphtalene, FLU: Fluorene, PHE: Phenanthrene, DBT: Dibenzothiophene, PYR: Pyrene, CHR: Chrysene*

*Dataset S2:* ***Phenanthrene metabolite tissue (ug PAH/g ww) content at all stages in early and late exposure.*** *PHE: Phenanthrene, OH1PHE: 1-hydroxyphenanthrene, OH3PHE: 3-hydroxyphenanthrene*, OH4PHE: 4-hydroxyphenanthrene, OH9PHE: 9-hydroxyphenanthrene.
